# Trajectory inference of epithelial-centered neighborhood profiles reconstructs a pseudo-temporal continuum in idiopathic pulmonary fibrosis

**DOI:** 10.64898/2026.06.15.732243

**Authors:** Satoshi Nakamura, Kazuya Tsubouchi, Yoshihiro Yamamoto, Tomotsugu Takano, Kosei Nakatsuru, Tomoyoshi Takenaka, Mikiko Hashisako, Yoshinao Oda, Isamu Okamoto

## Abstract

Idiopathic pulmonary fibrosis (IPF) is characterized by complex lung architecture and spatially heterogeneous remodeling, which have hindered integrated analysis of cell-intrinsic activity and intercellular communication during disease progression. Here we profiled six IPF lung specimens comprising more than 630,000 cells using the Xenium 5k panel and developed an epithelial-centered neighborhood profiling framework based on the local cellular composition around each epithelial cell. This approach captured fibrosis-associated variation in epithelial niches without requiring predefined histological regions. Pseudo-temporal continuum inference of these profiles reconstructed a continuous axis that reflected the spatial progression of fibrotic remodeling from relatively preserved alveolar regions to fibrotic and airway-like remodeled regions. Within this spatial dataset, we mapped coordinated changes in epithelial states, local microenvironments, epithelial intracellular pathway activities, and directional interactions with neighboring cell types along the same axis. Our findings provide a spatial framework that generates testable hypotheses for progressive epithelial niche remodeling in IPF.

## INTRODUCTION

Idiopathic pulmonary fibrosis (IPF) is a chronic, progressive, and ultimately fatal interstitial lung disease of unknown cause^1^. The prognosis remains poor, with median survival of approximately 2–5 years after diagnosis^1–3^. A deeper understanding of the mechanisms driving remodeling is therefore required to identify disease-modifying targets.

A major obstacle to understanding IPF pathogenesis is the spatial complexity of the IPF lung, which comprises regionally specialized epithelial, mesenchymal, endothelial, and immune niches. In IPF, this complexity is amplified by spatially heterogeneous remodeling that is difficult to reproduce in vitro or in experimental models^4,5^. Among the pathological changes in IPF, epithelial remodeling is particularly prominent^6–8^. Normal alveolar type 1 (AT1) and type 2 (AT2) cells are progressively lost, while *KRT5*^-^/*KRT17*^+^ aberrant epithelial cells emerge near fibroblastic foci^8–10^, and honeycomb cysts are lined by airway-like epithelium^11^. These observations suggest a directional transition from preserved alveolar epithelium to aberrant epithelial states and airway-like metaplasia. However, how this transition is coordinated with changes in the surrounding microenvironment remains incompletely understood.

Single-cell transcriptomics and spatial profiling have identified IPF-associated cell states and multicellular niches^7,10,12–15^. However, most spatial niche analyses describe tissue architecture as discrete structural states, leaving their relationship to fibrotic progression uncertain. Although airspace-based analyses have inferred fibrotic stages from individual airspace units, they provide limited access to cell-level transcriptional activity and cell–cell communication within the same spatial dataset^13^. Thus, a framework that links tissue architecture, epithelial remodeling, intracellular programs, and intercellular signaling within spatial data is needed.

Here, we analyzed six human IPF tissue samples using Xenium 5k spatial transcriptomics. We developed epithelial-centered neighborhood profiling to quantify the cellular composition surrounding each epithelial cell and used trajectory inference to reconstruct a pseudo-temporal continuum of fibrotic remodeling from local tissue architecture. We then integrated this continuum with cell-level pathway activity analysis and spatially resolved ligand–receptor scoring to examine the transition of epithelial intracellular programs and cell–cell communication within the same dataset. This framework provides a spatially anchored view of epithelial remodeling in IPF and identifies candidate signaling axes that may be associated with fibrotic progression.

## RESULTS

### Cell-type annotation and spatial organization of the IPF

We microdissected fibrotic regions with a histological usual interstitial pneumonia pattern from resected lung specimens from four patients with clinically and radiologically diagnosed idiopathic pulmonary fibrosis undergoing lung cancer surgery, yielding six tissue samples (Supplementary Table 1). The samples were profiled using the Xenium 5k platform (10x Genomics). The Xenium output was subsequently reprocessed with ProSeg^16^ to refine cell segmentation and reduce erroneous transcript assignment arising from ambient background signal or spillover from neighboring cells. After ProSeg-based processing and quality control, we retained 630,383 cells and 270,458,182 detected transcripts across 5,001 genes. The resulting cell-by-gene expression matrix was processed using standard a single-cell analysis workflow, including normalization, dimensionality reduction, and clustering (Fig. 1a). Major lineages were identified (Fig. 1b, Extended Data Fig. 1a, Supplementary Tables 2, 3) and then subclustered to resolve epithelial cells, fibroblasts, macrophages, endothelial cells, and pericytes/smooth-muscle cells (Fig. 1c–g, Extended Data Fig. 1b–f, Supplementary Tables 3–8). To refine annotations within the epithelial, fibroblast, and macrophage compartments, we imputed expression for genes not included in the 5k panel by integrating two publicly available single-cell RNA-seq reference datasets^7,14^ (Supplementary Fig. 1). Ultimately, we annotated 39 distinct cell types (Extended Data Fig. 2, Supplementary Table 9).

**Fig. 1.**
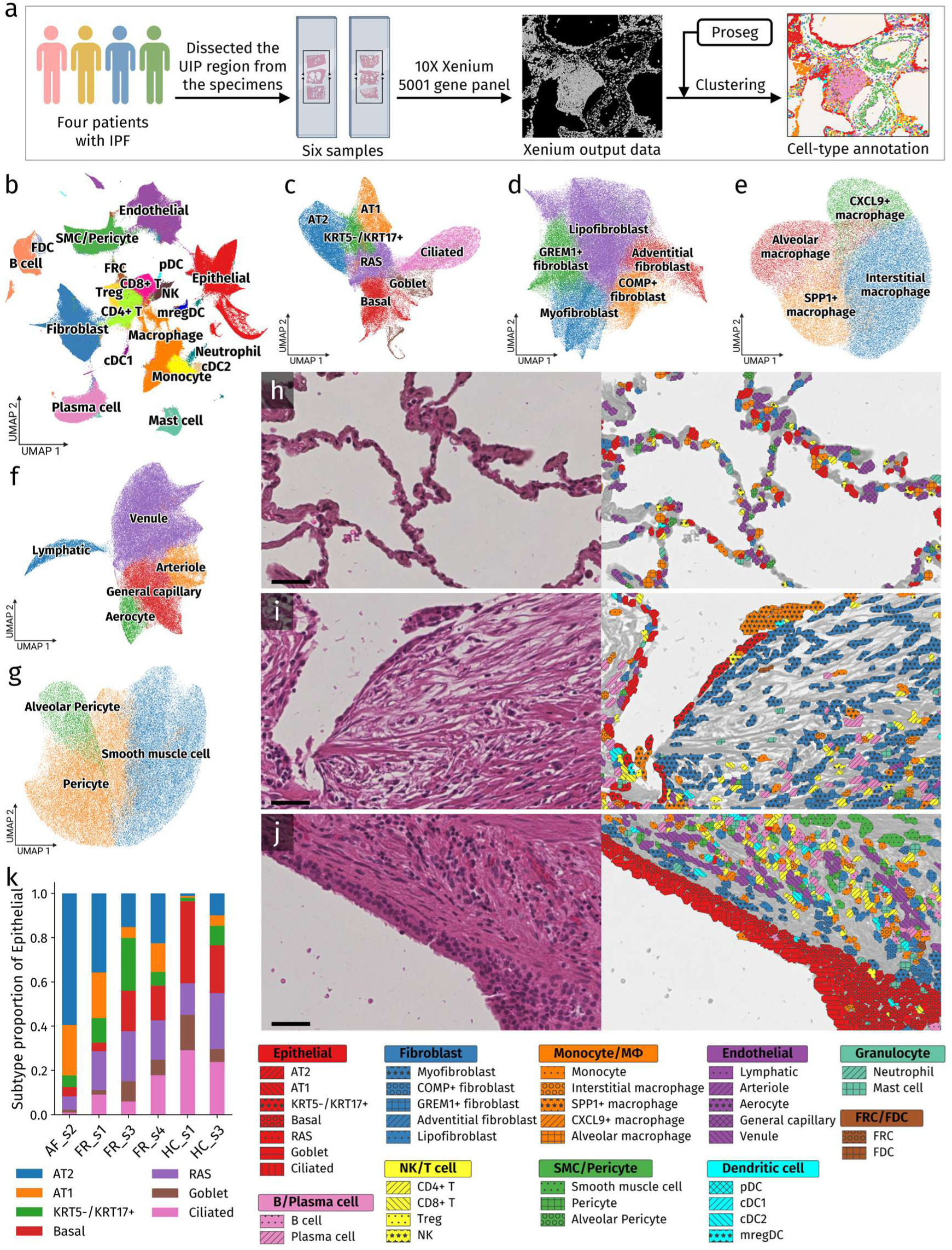
Cell-type and subtype annotation of IPF lung tissues using Xenium. **a**, Schematic overview of the study design and analysis workflow. Lung tissues with usual interstitial pneumonia morphology were microdissected from surgical specimens of patients with IPF, profiled using the Xenium 5k panel, processed with ProSeg-based segmentation, and annotated by clustering analysis. **b,** UMAP embedding of all cells colored by initial cell-type labels before lineage-specific subclustering and subtype refinement. **c–g,** UMAP embeddings of lineage-specific subclustering for epithelial cells (c), fibroblasts (d), macrophages (e), endothelial cells (f), and smooth muscle cells/pericytes (g). **h–j,** H&E images and corresponding Xenium/ProSeg cell-type segmentation maps showing preserved alveolar architecture (AF_s2; h), a fibroblastic focus (FR_s3; i), and airway epithelial metaplasia in honeycomb lung (HC_s1; j). Cell colors indicate annotated cell types as shown in the legend. Scale bars, 50 μm. **k,** Stacked bar plots showing epithelial subtype composition in each sample. DC, dendritic cell; pDC, plasmacytoid dendritic cell; cDC1, conventional dendritic cell type 1; cDC2, conventional dendritic cell type 2; mregDC, mature dendritic cell enriched in immunoregulatory molecules; FRC, fibroblastic reticular cell; FDC, follicular dendritic cell; SMC, smooth muscle cell; RAS, respiratory airway secretory; Mφ, macrophage.

To validate these annotations, we visualized cell-type distributions on the spatial maps (Extended Data Fig. 3, 4). Spatial inspection supported the annotations, showing preserved alveolar epithelial architecture (Fig. 1h), a fibroblastic focus in which *KRT5*^-^ /*KRT17*^+^ aberrant epithelial cells and myofibroblasts were in close proximity (Fig. 1i), consistent with prior reports describing their emergence during fibrotic progression^7,9,14^, and airway epithelial metaplasia in honeycomb lung (Fig. 1j). We also observed bronchial arteries and lymphatic vessels accompanying the airways with the bronchovascular bundles (Extended Data Fig. 5a, b), alveolar macrophages and *SPP1*^+^ macrophages extending into alveolar spaces (Extended Data Fig. 5c–e), and tertiary lymphoid structures within the lung interstitium (Extended Data Fig. 5c, f), all in expected anatomical or pathological contexts^13,17–19^. To examine the spatial organization of the annotated cell types, we performed neighborhood enrichment analysis to globally assess cell-type proximity across the tissue (Supplementary Fig. 2a). We observed spatial co-localization among alveolar unit components, including AT1/AT2, capillary endothelial cells, and aerocytes; among airway epithelial subtypes, including basal, respiratory airway secretory (RAS), goblet, and ciliated cells; and between *KRT5*^-^ /*KRT17*^+^ cells and myofibroblasts.

Aggregating cell-type counts across all samples, we detected substantial numbers of fibrosis-associated subtypes^7,14,20–22^, including *KRT5*^-^/*KRT17*^+^ aberrant epithelial cells, myofibroblasts, and *SPP1*^+^ macrophages (Supplementary Fig. 2b), supporting their inclusion in subsequent quantitative and statistical analyses. At the level of major cell types, proportions were broadly similar across samples, with no marked depletion or overrepresentation of any major lineage in a single specimen (Supplementary Fig. 3a), arguing against downstream results being driven primarily by sample-specific lineage imbalance. By contrast, subtype-level compositions within the epithelial, fibroblast, and endothelial compartments differed between samples (Fig. 1k, Supplementary Fig. 3b–e, Supplementary Table 10). Given that the samples were selected to capture distinct histopathologic stage of fibrosis, this variability likely reflects biological differences in fibrotic stage rather than instability in overall cell-type representation.

### Identifying spatial features of IPF through epithelial-centered neighborhood profiling

IPF is thought to be initiated by repetitive alveolar epithelial injury, followed by complex interactions between epithelial cells and surrounding mesenchymal, endothelial and immune cells that promote fibroblastic foci formation and, ultimately, bronchiolar metaplasia with honeycomb remodeling^1,10,23^. Previous niche-analysis frameworks have characterized IPF tissue architecture by analyzing spatial neighborhoods across all cells, leading to the identification of fibrosis-associated niches^13,15,24^. However, these global approaches treat all cells equivalently, and the resulting niches are therefore not necessarily anchored to epithelial remodeling states. We therefore hypothesized that epithelial-anchored niche profiling might better reflect microenvironmental changes accompanying fibrosis. To test this, we performed “epithelial-centered neighborhood profiling” by defining all cells within an 80-µm radius of each epithelial cell as its neighborhood and summarizing local cellular composition (Fig. 2a). Neighborhood sizes were broadly similar across samples and included adequate numbers of neighboring cells (Extended Data Fig. 6a). Clustering of the epithelial neighborhood composition matrix identified seven region-associated epithelial neighborhood classes: near-normal alveolar, pre-fibrotic, fibrotic, metaplastic, bronchiolar, alveolar duct, and epithelial-rich (Fig. 2b).

**Fig. 2.**
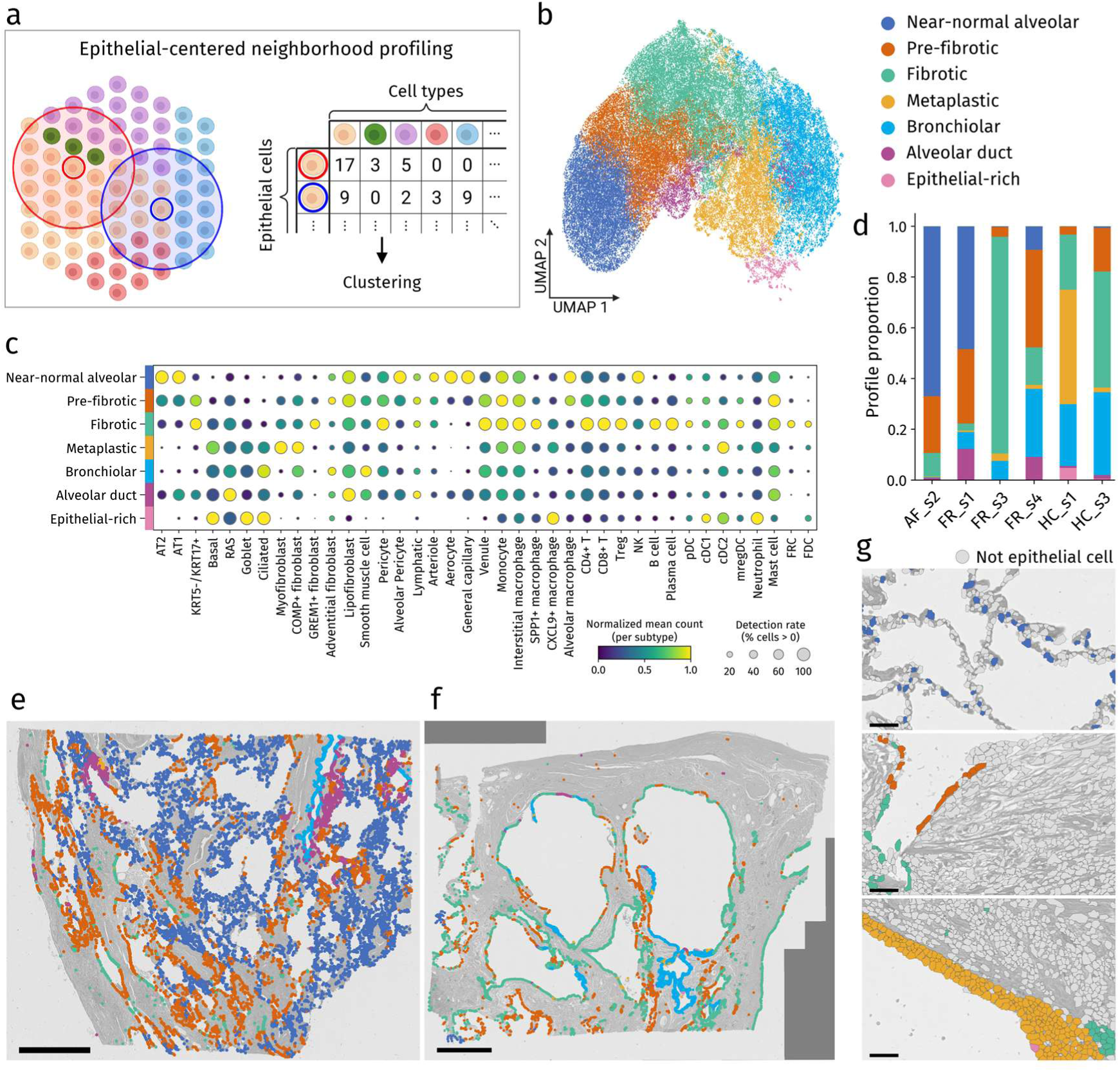
Epithelial-centered neighborhood profiling identifies spatially distinct neighborhood classes in IPF. **a**, Schematic overview of epithelial-centered neighborhood profiling. For each epithelial cell, neighboring cells within an 80-µm radius were counted by annotated cell type to generate an epithelial cell-centered neighborhood profile, which was then used for clustering. **b,** UMAP embedding of epithelial-centered neighborhood profiles. Seven neighborhood classes were identified based on neighboring cell-type composition, central epithelial identity, and spatial localization in tissue. **c,** Dot plot showing the normalized mean count of each cell type or subtype within each neighborhood class. Dot size indicates the detection rate, defined as the percentage of epithelial-centered profiles in which each cell type or subtype was detected. **d,** Stacked bar plots showing the composition of neighborhood classes in each sample. **e, f,** Spatial maps of epithelial cells color-coded by neighborhood class overlaid on grayscale tissue images in FR_s1 (e) and HC_s3 (f). **g,** Magnified spatial maps of epithelial cells color-coded by neighborhood class in regions corresponding to preserved alveolar architecture (top; same region as Fig. 1h), a fibroblastic focus (middle; same region as Fig. 1i), and airway epithelial metaplasia in honeycomb lung (bottom; same region as Fig. 1j). Non-epithelial cells are shown in gray. Scale bars, 1,000 μm (e, f) and 50 μm (g).

Examination of epithelial neighborhood composition revealed distinct cellular microenvironments (Fig. 2c). The near-normal alveolar class was enriched for AT1/AT2 cells together with alveolar pericytes, aerocytes, general capillary endothelial cells, and alveolar macrophages. Pre-fibrotic and fibrotic classes shared enrichment of *KRT5^−^/KRT17^+^*aberrant epithelial cells, myofibroblasts, *GREM1^+^* fibroblasts, and diverse immune populations, but differed in the balance between AT1/AT2 cells and *KRT5^−^/KRT17^+^* cells: the pre-fibrotic class retained higher proportions of AT1/AT2 cells, whereas the fibrotic class showed greater enrichment of *KRT5^−^/KRT17^+^* cells and *GREM1^+^* fibroblasts, which motivated the pre-fibrotic and fibrotic annotations. The metaplastic class was characterized by airway epithelial cells accompanied by myofibroblasts and *COMP*^+^ fibroblasts. The bronchiolar class also contained abundant airway epithelial populations but showed fewer myofibroblasts and enrichment of adventitial fibroblasts and smooth-muscle cells. Some bronchiolar-class profiles were located in airway-remodeled regions on tissue sections, suggesting partial overlap with metaplastic epithelial remodeling. We also identified an alveolar duct class marked by a RAS-rich epithelial component. Finally, we identified an epithelial-rich class composed almost exclusively of airway epithelial cells.

Class abundance varied across samples in a manner consistent with histopathology (Fig. 2d). For instance, AF_s2, a boundary region between preserved alveoli and fibrotic remodeling, was dominated by near-normal alveolar and pre-fibrotic classes, whereas HC_s1, derived from honeycomb regions, was enriched for fibrotic and metaplastic classes. Spatial mapping showed intermingled small-scale patches, highlighting the pronounced spatial heterogeneity of fibrotic remodeling in IPF (Fig. 2e, f, Extended Data Fig. 6b–e). These neighborhood classes also matched representative preserved alveoli, fibroblastic foci and honeycomb lung in Fig. 1h–j (Fig. 2g). The bronchiolar class mapped to airway regions, and the alveolar duct class was adjacent to near-normal alveolar areas (Extended Data Fig. 5a, 6f, g). As alveolar ducts are RAS-enriched structures that connect terminal bronchioles to alveoli^25^, the emergence of the alveolar duct class supported the anatomical validity of the profiling framework (Extended Data Fig. 6g). The epithelial-rich class localized to the luminal or apical side of expanded airway epithelium, suggestive of epithelial hyperplasia and detached epithelial aggregates (Fig. 2f, Extended Data Fig. 6h). Together, epithelial-centered neighborhood profiling distinguished fibrosis-associated contexts from non-fibrotic ones and captured phase-like microenvironmental differences during remodeling.

### An epithelial-centered pseudo-temporal continuum resolves staged remodeling in IPF

Given that fibrotic remodeling in IPF proceeds unidirectionally within the local epithelial microenvironment, yet distinct histological stages coexist within the same specimen, we reasoned that epithelial-centered neighborhood profiles could be ordered along a pseudo-temporal continuum of fibrotic progression. We adapted the logic of pseudotime analysis, which infers dynamic biological processes from snapshot single-cell data containing mixed cellular states. Here, instead of gene expression, we used cell-type composition vectors derived from each epithelial-centered neighborhood profile as input, allowing continuity to be inferred from differences in local cellular composition. As the input and interpretation differ from conventional pseudotime analysis, we refer to this framework as “pseudo-temporal continuum inference” (Fig.3a). To focus on fibrotic remodeling, we excluded bronchiolar and alveolar duct classes, which reflected bronchiolar and alveolar duct structures, and the epithelial-rich class, which contained little non-epithelial microenvironmental information. Using near-normal alveolar, pre-fibrotic, fibrotic, and metaplastic classes, we inferred trajectory structure with scFates^26^ and designated profiles assigned to the near-normal alveolar class as the root to assign directionality (Fig. 3b, Extended Data Fig. 7a). Each epithelial cell was assigned a continuum position (Extended Data Fig. 7b), and projection of these values onto tissue coordinates revealed a spatially continuous distribution of fibrotic remodeling across samples (Fig. 3c, d, Extended Data Fig. 7c–f). Importantly, when samples were ranked by mean continuum position, they followed the histopathological severity order from AF through FR to HC, supporting the ability of this continuum-based framework to capture the pathological progression across samples (Fig. 3e).

**Fig. 3.**
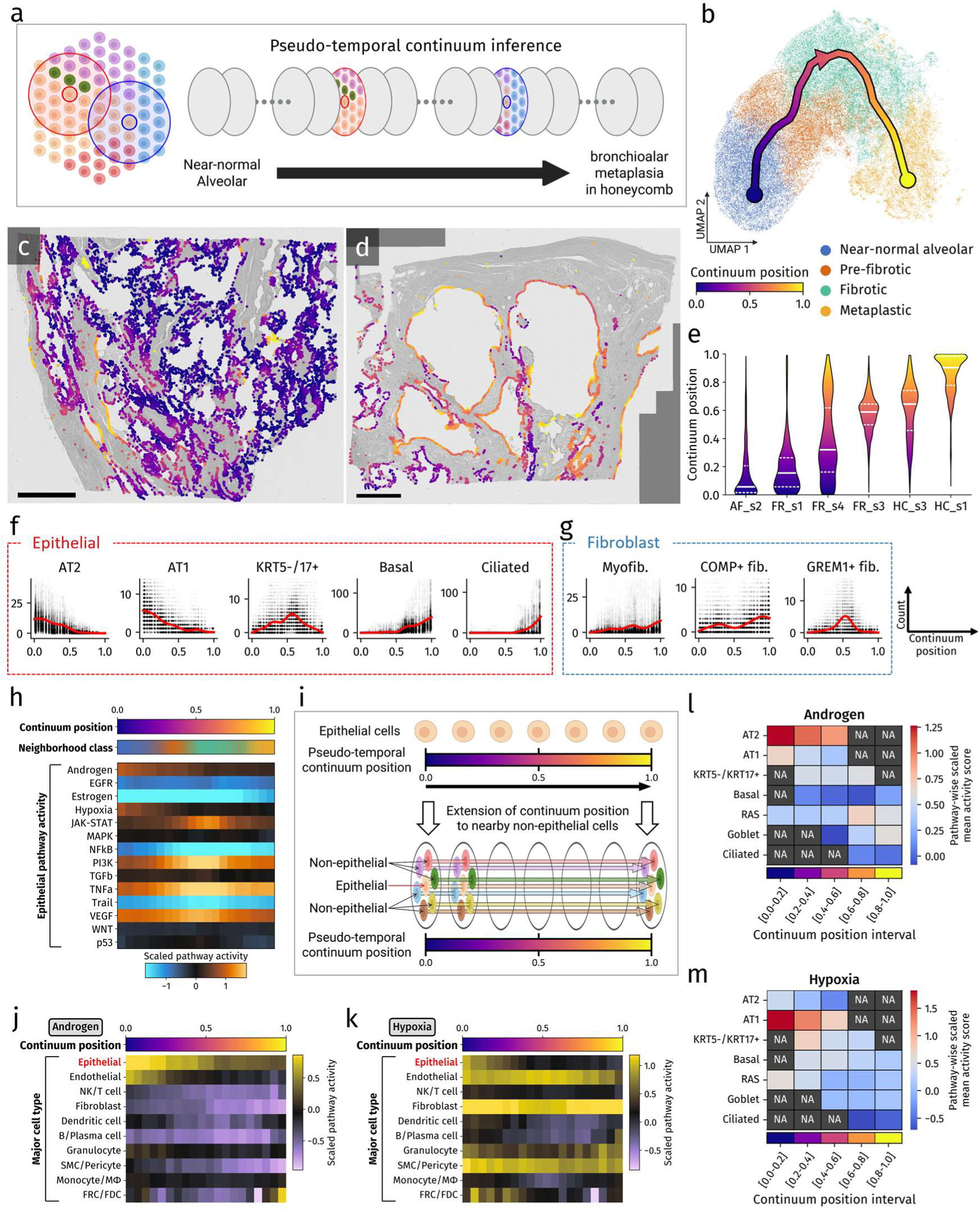
Pseudo-temporal continuum inference from epithelial-centered neighborhood profiles reveals dynamic epithelial and microenvironmental programs. **a**, Schematic overview of pseudo-temporal continuum inference based on epithelial-centered neighborhood profiles. Cell-type composition vectors around individual epithelial cells were used as input to infer a continuum of fibrotic remodeling from near-normal alveolar regions toward bronchiolar metaplasia in honeycomb lung. **b,** UMAP embedding of epithelial-centered neighborhood profiles with the inferred continuum trajectory. Bronchiolar, alveolar duct, and epithelial-rich neighborhood classes were excluded from this analysis to focus on fibrotic remodeling rather than normal airway-associated structures or epithelial-dominant profiles with limited non-epithelial neighborhood information. **c, d,** Spatial maps of epithelial cells color-coded by continuum position overlaid on grayscale tissue images in FR_s1 (c) and HC_s3 (d). Scale bars, 1,000 μm. **e,** Violin plots showing the distribution of continuum positions across samples. Solid horizontal lines indicate the mean, and dashed lines indicate the first and third quartiles. Samples are ordered by increasing mean continuum position. **f, g,** Changes in the numbers of selected epithelial subtypes (f) and fibroblast subtypes (g) within epithelial-centered neighborhood profiles along the continuum. Gray dots indicate observed neighborhood counts, and red curves indicate fitted trends inferred by scFates. **h,** Heatmap showing epithelial intracellular pathway activity inferred using PROGENy across 20 equal-width bins of continuum position. Pathway activity scores were averaged within each bin and scaled by pathway. **i,** Schematic overview of extension of continuum position to nearby non-epithelial cells. Non-epithelial cells located within 30 μm of epithelial cells were assigned local continuum positions, enabling analysis of microenvironmental cell states along the epithelial remodeling continuum. **j, k,** Heatmaps showing Androgen (j) and Hypoxia (k) pathway activity in epithelial and nearby non-epithelial major cell types along the continuum. Activity scores were averaged within continuum-position bins and scaled for visualization. **l, m,** Heatmaps showing subtype-resolved epithelial Androgen (l) and Hypoxia (m) pathway activity across five continuum-position intervals. Mean pathway activity was calculated for each epithelial subtype within each interval. Intervals in which a subtype accounted for less than 3% of epithelial cells were excluded and are shown as NA.

To assess whether the inferred continuum recapitulated the established sequence of epithelial remodeling in IPF and whether this was accompanied by coordinated changes in the surrounding microenvironment, we examined how epithelial and non-epithelial cell populations varied along the continuum. Each profile retained counts for all surrounding cell types, enabling us to track cell-type composition changes across the local epithelial microenvironment (Fig. 3f, g, Supplementary Fig. 4). AT1/AT2 cells were most abundant at the beginning of the continuum, declined progressively through the early-to-intermediate phase (continuum positions 0.0–0.5), and remained low thereafter. *KRT5^−^/KRT17^+^* cells increased across the same interval and then decreased. In contrast, airway epithelial populations emerged from the intermediate phase onward and increased progressively toward the late phase (continuum positions 0.5–1.0; Fig. 3f). These changes recapitulated the reported transition from alveolar epithelium to aberrant epithelial states and ultimately to bronchiolar metaplasia in IPF^7,14,27^. Surrounding non-epithelial populations also underwent coordinated remodeling along this continuum. Among fibroblasts, myofibroblasts and *COMP^+^* fibroblasts increased progressively across the continuum, whereas *GREM1^+^* fibroblasts were enriched selectively in the intermediate phase (Fig. 3g).

We next examined changes in PROGENy^28^ pathway activity in epithelial cells along the pseudo-temporal continuum using decoupler^29^. Androgen and Hypoxia pathway activities decreased from the early to intermediate phase, while JAK–STAT, PI3K and VEGF activities peaked in the intermediate phase, TNFα activity remained positive throughout the continuum but was relatively attenuated in the early phase, and NF-κB activity remained negative throughout, with weaker suppression in the early phase. The remaining pathways showed little change across the continuum (Fig. 3h). To assess whether inferred epithelial pathway activity patterns could be influenced by transcript spillover from neighboring cells, we assigned each epithelial cell’s continuum position to all cells within a 30-μm radius, generating an extended continuum position that enabled enrichment analysis in non-epithelial populations (Fig. 3i). Under this framework, Androgen and Hypoxia showed activity patterns within epithelial cells that were distinct from those of neighboring cell types, supporting these changes as epithelial-specific (Fig. 3j, k). Subtype-resolved analysis across five continuum bins further showed that Androgen activity was highest in AT2 cells (Fig. 3l) and Hypoxia activity was highest in AT1 cells (Fig. 3m), and that both progressively declined along the continuum, even as AT2 and AT1 abundance decreased in parallel with the known epithelial remodeling trajectory in IPF (Extended Data Fig. 7g). Together, these findings indicate that the decreases in Androgen and Hypoxia activity reflect not only shifts in epithelial subtype composition but also attenuation of intracellular pathway activity within epithelial cells. In contrast, JAK–STAT, PI3K, VEGF, TNFα and NF-κB activities in epithelial cells showed temporal patterns similar to those observed in surrounding cell populations, with higher peak scores in the latter than in epithelial cells, suggesting that the corresponding epithelial pathway activity patterns may have been partly influenced by transcript spillover, although genuine intracellular pathway activity within epithelial cells could not be excluded (Extended Data Fig. 7h–l).

### Epithelial-centered cell–cell interaction analysis identifies subtype-specific signaling programs and highlights PERIOSTIN interactions in IPF lung

Having mapped the pseudo-temporal dynamics of epithelial neighborhoods an epithelial intracellular activities, we next evaluated cell–cell communication within the epithelial-centered microenvironment. We referenced known ligand–receptor interactions in the CellChat^30^ database and restricted the analysis to interactions for which all genes encoding the ligand and receptor components were included in the Xenium 5k gene panel (Supplementary Table 10). We then established an epithelial-centered, cell-level interaction scoring framework that quantified putative communication for each individual epithelial cell and its surrounding cells on the basis of ligand and receptor expression together with distance-based weighting between cells. Using this framework, we quantified bidirectional communication between epithelial cells and neighboring cells within 300 µm, including signals from epithelial cells to surrounding cells and those directed back toward epithelial cells (Fig. 4a; see Supplementary Note). We next aggregated the inferred interaction scores by cell subtype and visualized the top 50 significant interactions in Sankey diagrams (Fig. 4b, c). For interactions directed from neighboring cells to epithelial cells, *KRT5^−^/KRT17^+^* cells received prominent inputs from myofibroblasts and *SPP1^+^* macrophages, whereas AT2 cells were characterized by inputs from lipofibroblasts and alveolar macrophages (Fig. 4b). In the opposite direction, *KRT5^−^/KRT17^+^*cells showed a distinct outgoing interaction pattern directed toward immune cell populations as well as lipofibroblast and endothelial populations (Fig. 4c). Among these incoming interactions, signaling from myofibroblasts to *KRT5^−^/KRT17^+^*cells was among the strongest and was consistently dominated by PERIOSTIN-family interactions, primarily through POSTN–ITGAV_ITGB3 and POSTN–ITGAV_ITGB5 (Fig. 4d).

**Fig. 4.**
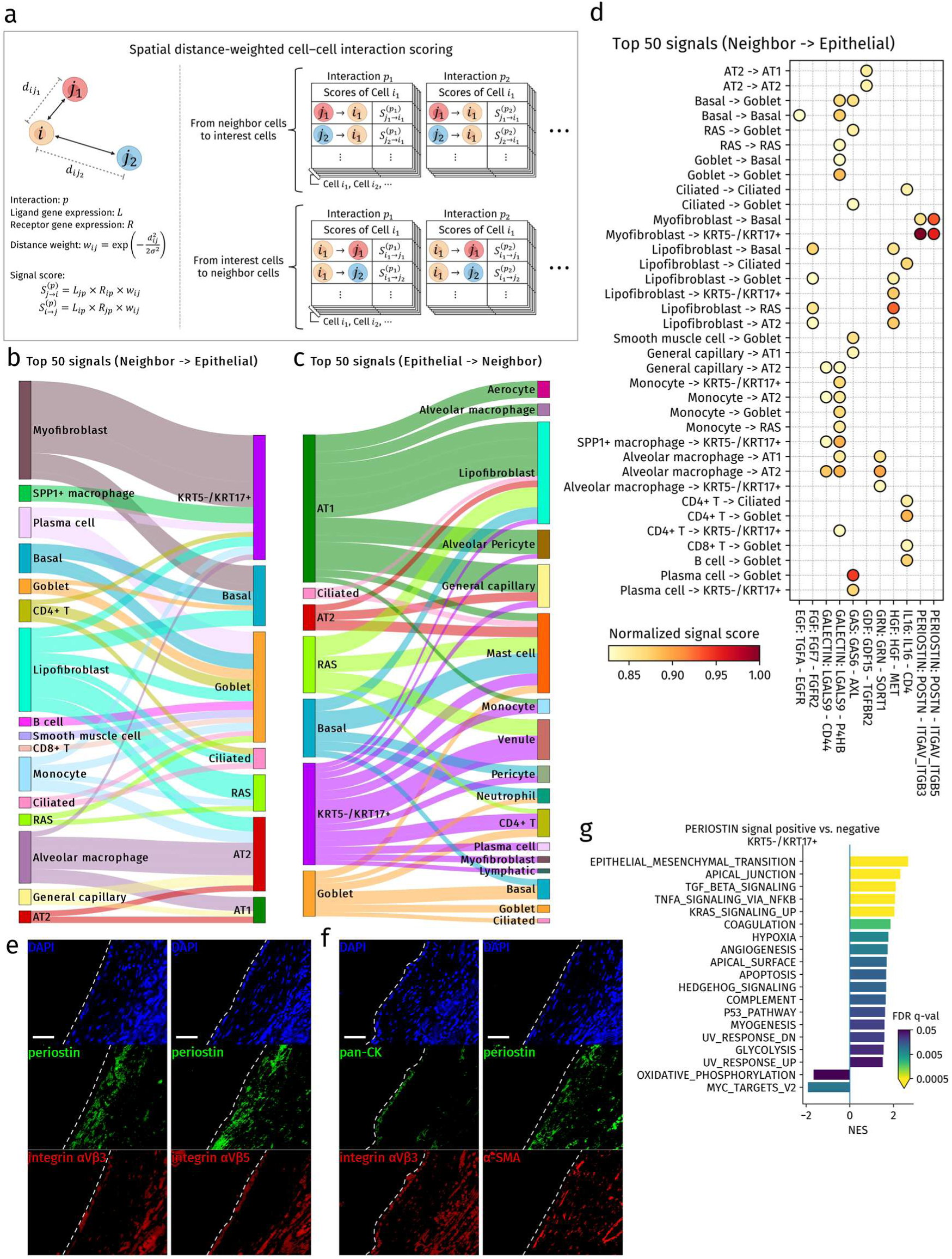
Spatial distance-weighted cell–cell interaction analysis identifies prominent periostin-family signaling in fibroblastic foci. **a**, Schematic overview of spatial distance-weighted cell–cell interaction scoring. For each ligand–receptor pair, interaction scores were calculated by combining ligand expression in sender cells, receptor expression in receiver cells, and a Gaussian distance-based weight. Scores were computed for interactions from neighboring cells to epithelial cells and from epithelial cells to neighboring cells within a 300-μm radius. **b, c,** Sankey diagrams summarizing the top 50 subtype-level interactions from neighboring cells to epithelial cell subtypes (b) and from epithelial cell subtypes to neighboring cells (c). **d,** Dot plot showing representative top interactions from neighboring cells to epithelial cell subtypes. Rows indicate sender cell types and receiver epithelial subtypes, and columns indicate ligand–receptor pairs. Dot color represents the normalized interaction score. **e,** Representative immunofluorescence images of fibroblastic foci in serial sections from FR_s1, showing DAPI (blue), periostin (green), and integrin αVβ3 or integrin αVβ5 (red). Scale bars, 50 μm. **f,** Representative immunofluorescence images of fibroblastic foci in serial sections from FR_s1, showing pan-CK, periostin, integrin αVβ3, and α-SMA. Co-staining showed periostin in α-SMA-positive myofibroblast-rich regions and integrin αVβ3 in pan-CK-positive epithelium overlying fibroblastic foci. Scale bars, 50 μm. **g,** Gene set enrichment analysis of *KRT5^−^/KRT17^+^* epithelial cells comparing cells positive versus negative for inferred incoming PERIOSTIN-family signals. Bars show normalized enrichment scores (NES) for significantly enriched Hallmark gene sets (FDR q value < 0.05), with color indicating the FDR q value.

The inferred ligand–receptor interactions were based on transcript-level data and could potentially be influenced by transcript spillover. We therefore sought protein-level support for the top-ranked PERIOSTIN pathway interactions. Immunofluorescence staining on serial sections from the same samples used for Xenium analysis revealed periostin-positive cells within fibroblastic foci in close proximity to overlying cells expressing integrin αVβ3 or αVβ5, supporting the presence of cell–cell interactions at the protein level (Fig. 4e, Extended Data Fig. 8a). Additional co-staining with α-SMA and pan-cytokeratin further showed that periostin-positive cells within fibroblastic foci were α-SMA-positive myofibroblasts, whereas integrin αVβ3-positive overlying cells were pan-cytokeratin-positive epithelial cells (Fig. 4f, Extended Data Fig. 8b). Together, these findings supported the biological plausibility of the inferred periostin–integrin interactions.

Building on the cell-level interaction scores assigned to individual epithelial cells, we examined whether these scores could be used to compare transcriptional programs associated with inferred incoming signals. For each interaction, epithelial cells were divided into signal-negativeand signal-positive groups according to whether the incoming interaction score was 0 or greater than 0. As this cell-level positive-versus-negative comparison represents a downstream extension of conventional ligand-receptor analysis, we asked whether it could recover known biological responses. We first examined TGFβ signaling, whose epithelial-mesenchymal transition (EMT)-promoting effects are well established across epithelial contexts^31,32^. For all epithelial cells combined, we generated sample-level pseudobulk profiles for signal-positive and signal-negative groups and compared them using decoupler (Extended Data Fig. 8c). To avoid circularity, ligand- and receptor-encoding genes used to define the interaction scores were excluded from the 5,001-gene panel before gene sets enrichment analysis (GSEA). The TGFβ signal-positive group showed enrichment of the EMT hallmark signature overall and in most epithelial subtypes (Extended Data Fig. 8d, e). We next examined HGF–MET signaling, which can promote EMT in cancer cells and other epithelial contexts but has also been reported to suppress EMT in alveolar epithelium^33,34^. EMT was negatively enriched in alveolar-associated epithelial subtypes, including AT2, AT1, and RAS cells, whereas it was positively enriched in basal and *KRT5^−^/KRT17^+^* cells (Extended Data Fig. 8f, g). Although these analyses do not prove *in vivo* signaling effects, recovery of expected context-dependent patterns supports this cell-level comparison strategy. The PERIOSTIN pathway showed the highest interaction scores in our analysis, and we therefore examined transcriptional effects in the same manner. PERIOSTIN signal-positive cells were predominantly found in the *KRT5^−^/KRT17^+^* and basal cells (Extended Data Fig. 8h), both of which mainly received input from myofibroblasts (Fig. 4d). In these subtypes, EMT, TGFβ signaling, and TNFα signaling were commonly enriched, suggesting a shared transcriptional program associated with myofibroblast-derived PERIOSTIN-family interactions (Fig. 4g, Extended Data Fig. 8i).

### Epithelial cell**–**cell communication is dynamically remodeled along the fibrotic continuum

Our cell–cell interaction analysis provided a snapshot view that did not account for temporal progression. We therefore assessed how specific interactions changed along the pseudo-temporal continuum by binning continuum position into equal-width intervals and re-aggregating epithelial interaction scores within each bin (Fig. 5a, b, Supplementary Tables 13, 14). In the neighbor-to-epithelial direction, individual interactions within the EGF, TGFβ, and FGF families were enhanced at different phases along the continuum (Fig. 5a). For example, within the TGFβ pathway, interactions converging on ACVR1B_TGFbR2 showed different temporal patterns, reflecting differences in ligand-producing cell types and timing between TGFB1-, TGFB3-, and TGFB2-mediated interactions (Extended Data Fig. 9a-c). POSTN–ITGAV_ITGB3 and POSTN–ITGAV_ITGB5 both showed high scores at the intermediate stage, where fibrotic remodeling appeared most active (Fig. 5c, Extended Data Fig. 9d).

**Fig. 5.**
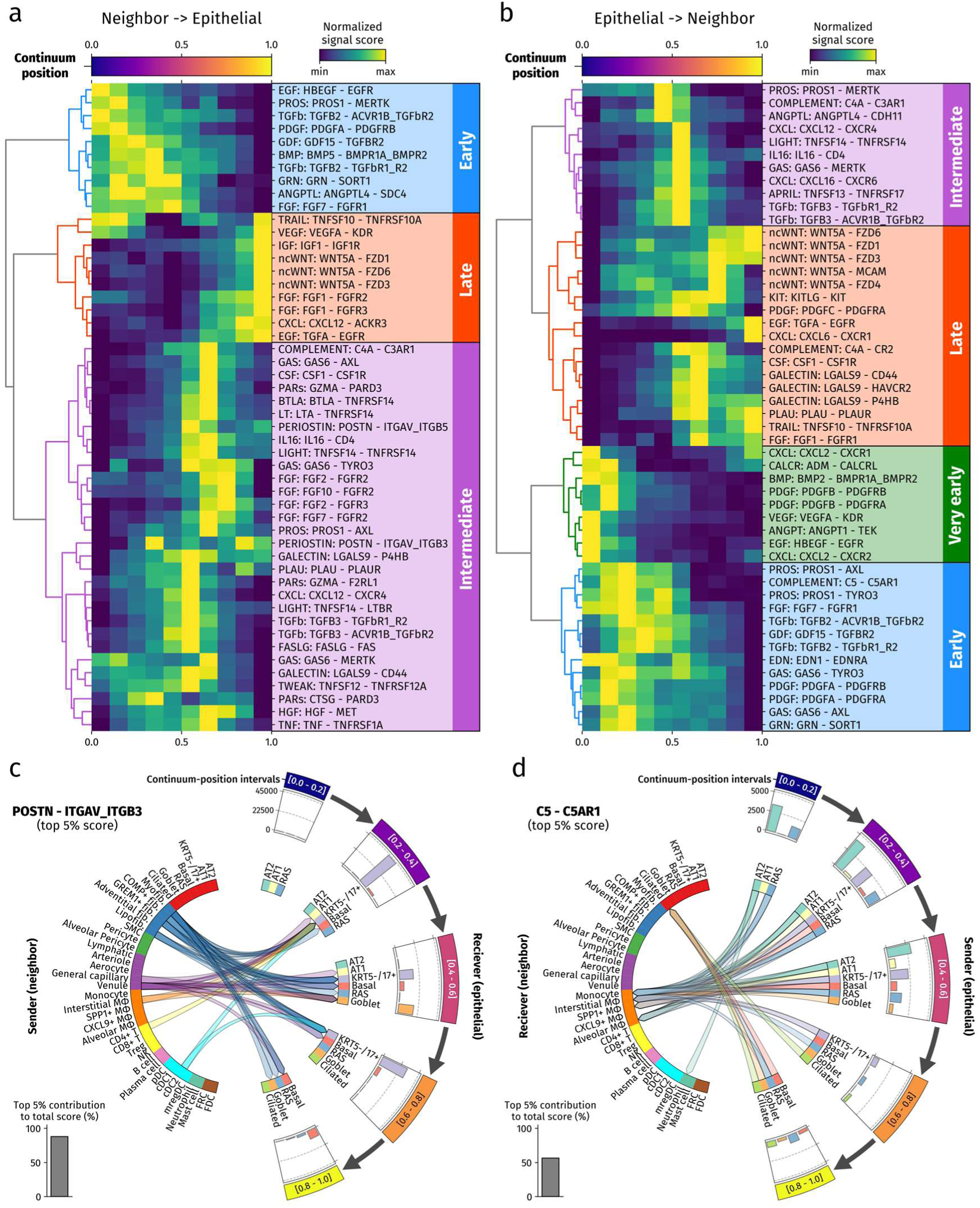
Dynamic changes in cell–cell interactions along the pseudo-temporal continuum. **a, b**, Heatmaps showing the dynamics of cell–cell interaction scores along the continuum for interactions from neighboring cells to epithelial cells (a) and from epithelial cells to neighboring cells (b). The continuum was divided into 10 bins, and interaction scores were aggregated within each bin and normalized by the number of epithelial cells in that bin. The top 50 interactions ranked by total score across bins are shown. Rows were hierarchically clustered according to their temporal patterns, and major activation patterns are annotated at right. Color intensity indicates the interaction score scaled within each interaction. **c, d,** Circular plots showing the distribution of representative interactions across cell subtypes and continuum intervals for POSTN–ITGAV_ITGB3 from neighboring cells to epithelial cells (c) and C5–C5AR1 from epithelial cells to neighboring cells (d). Epithelial cells were grouped into five continuum-position intervals. Epithelial subtype–interval combinations in which the subtype accounted for less than 3% of epithelial cells in that interval were excluded from visualization. Links indicate interactions between sender and receiver cell subtypes, and only interactions belonging to the top 5% of scores are shown. Bar plots above the epithelial sectors show the summed interaction score within each continuum-position interval. The inset bar plot indicates the percentage contribution of the top 5% of interaction scores to the total score.

We next examined signals directed from epithelial cells to neighbor cells and found that several pathways strongly linked to fibrosis, including VEGF, PDGF, and TGFβ, were already evident at an early stage of the continuum (Fig. 5b). More specifically, VEGFA–KDR signaling was largely restricted to AT1 cells and targeted cell types associated with normal alveolar architecture, including alveolar pericytes, aerocytes, and lipofibroblasts (Extended Data Fig. 10a). This pattern is consistent with a homeostasis-associated role for VEGFA–KDR signaling within the alveolar niche^35,36^. In contrast, PDGF-related interactions, although also partly derived from AT1 cells, showed high signal scores from *KRT5^−^/KRT17^+^*and basal cells and were directed toward cell types that increased during fibrosis, including myofibroblasts, *GREM1^+^* fibroblasts, and venules (Extended Data Fig. 10b, c). These findings suggest that epithelial PDGF signaling may contribute to pro-fibrotic remodeling during disease progression. Finally, we examined signaling interactions enriched at the early stage of the continuum and selected the C5–C5AR1 axis as a representative example for further characterization (Fig. 5d). This interaction was observed from the earliest continuum bins and was driven predominantly by signaling from AT2 cells toward monocytes and macrophages. We then compared AT2 cells with positive versus negative epithelial-to-neighbor C5–C5AR1 interaction scores (Extended Data Fig. 10d). C5–C5AR1-positive AT2 cells showed enrichment of E2F targets, glycolysis, and oxidative stress-related pathways (Extended Data Fig. 11e). These transcriptional features were suggestive of a distinct epithelial state characterized by cell-cycle engagement, metabolic rewiring, and oxidative stress.

## DISCUSSION

In this study, we analyzed six IPF tissue samples using Xenium 5k panel-based spatial transcriptomics and developed an epithelial-centered framework to characterize fibrotic remodeling. Pseudo-temporal continuum inference based on epithelial-centered neighborhood profiling allowed us to model fibrotic remodeling as a continuous process anchored to the local cellular environment of epithelial cells. This continuum reflected histopathologic progression and provided a common axis for evaluating epithelial subtype composition, neighboring cell populations, epithelial intracellular pathway activity, and inferred interactions between epithelial cells and their surrounding cells during fibrotic progression. Although the findings should be interpreted as hypothesis-generating, they provide a spatially anchored framework for understanding the multicellular coordination of fibrotic remodeling in IPF.

Previous spatial and single-cell studies of pulmonary fibrosis have often defined disease progression at the sample or region level, and even niche-based analyses typically considered all cell types together, resulting in stage-like rather than continuous representations of remodeling^12,13,15,24^. By reducing the spatial heterogeneity of IPF into epithelial-centered neighborhood profiles representing different phases of fibrotic remodeling, our approach incorporated continuity into the analysis of epithelial microenvironments. This framework enabled pathway activity and cell–cell interactions to be evaluated within the same spatial dataset along a shared remodeling axis, without relying on separate datasets or region-level annotations. In addition, retaining inferred interaction scores at the single-cell level allowed signal-positive and signal-negative epithelial cells to be compared within tissue, providing an observational proxy of how epithelial states differ according to local inferred signaling input or output. Integrating these cell-level interaction scores with the pseudo-temporal continuum further allowed us to examine how epithelial–neighbor interactions varied along the fibrotic remodeling trajectory.

Analysis of epithelial intracellular pathway activity suggested that Androgen and Hypoxia pathway activities decreased along the continuum. Among the pathway changes observed along the continuum, the decline in Androgen pathway activity was particularly relevant because androgen receptor expression has been reported in normal alveolar epithelial cells^37^, and androgen-related signaling has been implicated in epithelial proliferation and repair^38^. Epidemiologic observations that male sex is a risk factor for IPF, whereas higher serum testosterone levels are associated with lower IPF risk^1,39,40^, suggest that reduced androgen-related signaling may be relevant to IPF development despite the male predominance of the disease. These findings raise the possibility that attenuation of androgen-related epithelial programs may be associated with impaired alveolar epithelial homeostasis during early fibrotic remodeling. Hypoxia pathway activity was enriched in early AT1-associated alveolar states but declined along the continuum. Given links between hypoxia, hypoxia-inducible factor signaling, alveolar epithelial injury and profibrotic responses^41,42^, this pattern suggests that hypoxia-related epithelial programs may be involved in early alveolar remodeling but become less prominent later.

Among the inferred interactions, periostin-related signaling from myofibroblasts to *KRT5^−^/KRT17^+^* and basal cells was particularly prominent. Periostin has been implicated in IPF as a profibrotic matricellular protein and a biomarker of disease progression^43,44^, and it can interact with TGFβ signaling in lung fibroblasts through integrin αVβ3/αVβ5 and Smad3-dependent mechanisms^45^. Consistent with these observations, epithelial cells with positive periostin-related interaction scores showed enrichment of EMT-, TGFβ-, and TNFα-related programs. Periostin-related interactions were also enriched in the intermediate phase of the continuum, during which *KRT5^−^/KRT17^+^*aberrant epithelial cells became prominent. These findings suggest that periostin-related signaling may represent a profibrotic mesenchymal–epithelial interaction axis. This axis is not readily captured by global cell–cell interaction analyses but can be resolved by focusing on epithelial-centered neighborhoods.

This study has several limitations. First, the sample size was small and the study was primarily computational, so inter-individual variability and sampling bias may have affected the results. The continuum was constructed at single-cell resolution and was not restricted to a single sample, but the findings require validation in larger independent cohorts and experimental systems. Second, the Xenium 5k panel limited pathway inference and ligand–receptor analysis; interactions involving genes outside the panel could not be evaluated. Third, segmentation errors and transcript spillover remain important technical concerns, particularly for signal-positive versus signal-negative comparisons, where inferred signaling may reflect local proximity as well as true biological communication.

In summary, this study presents an epithelial-centered spatial transcriptomic framework for reconstructing a pseudo-temporal continuum of fibrotic remodeling. By linking this continuum to epithelial pathway activity and cell-level interaction modeling, we identified candidate epithelial programs and intercellular signaling axes associated with IPF progression. Although these findings remain observational, this framework may provide a basis for future studies of epithelial–microenvironment remodeling in IPF.

## METHODS

### Sample preparation

Lung resection was performed as therapeutic management for lung cancer associated with IPF at Kyushu University Hospital. Resected specimens were processed as formalin-fixed, paraffin-embedded (FFPE) tissue blocks. From these FFPE specimens, we retrospectively selected four patients whose fibrotic portion was pathologically diagnosed as showing a usual interstitial pneumonia (UIP) pattern. Clinical characteristics, surgical procedures, and resection sites are summarized in Supplementary Table 1. For each block, fibrotic areas were identified by pathologists and respiratory physicians. After review of the entire resected specimen, we microdissected regions representing a spectrum of fibrotic remodeling, including relatively preserved alveolar architecture, fibroblast focus-rich areas, and honeycombing, for subsequent Xenium profiling. Based on histopathologic assessment, samples were categorized as fibroblastic focus-rich (FR_s1, FR_s3, FR_s4), honeycombing-dominant (HC_s1, HC_s3), or representing the interface between relatively preserved alveolar architecture and fibrotic remodeling (AF_s2). This study was performed in accordance with the Declaration of Helsinki and was approved by the Institutional Review Board of Kyushu University Hospital (approval numbers 23495-00). The requirement for written informed consent was waived, and an opt-out method was used because of the retrospective nature of the study.

### Xenium in situ workflow

We performed in situ transcriptomic profiling using the Xenium in situ platform (10x Genomics) with the Xenium Prime 5K Human Pan Tissue & Pathways Panel (PN-1000724). From the FFPE tissue blocks, six samples derived from UIP regions were sectioned and mounted on Xenium slides (three samples per slide, two slides in total). FFPE sections were processed following the manufacturer’s demonstrated protocol for deparaffinization and decrosslinking (CG000580). Subsequent steps, including probe hybridization, ligation, rolling-circle amplification, and iterative fluorescent probe hybridization with imaging, were carried out on the Xenium Analyzer according to the standard workflow. Xenium output data was processed with Xenium ranger v3.1.0.0 and contained transcript coordinates with quality value (Q-score) ≥20 and cell membrane and nucleus segmentation information generated from default staining.

### Re-segmentation with ProSeg

Xenium Onboard Analysis output was re-segmented using ProSeg^16^. As input, we provided the Xenium transcript table, which contains per-transcript gene identity, 3D coordinates in microns, the Xenium-provided preliminary cell assignment, and a nuclear overlap indicator. We used the Xenium preset and default parameters without manual tuning. The resulting cell polygons and cell-by-gene count matrix were used for all downstream analyses.

### Quality control

For each Xenium dataset, we performed per-cell barcode filtering to remove low-quality cells before downstream analyses. We calculated, per cell, the ratio of high-quality (Q-score ≥20) transcript counts to total molecular counts and excluded cells with a ratio <0.95. To further remove extremely sparse profiles, we retained cells with ≥15 detected transcripts. These thresholds were applied uniformly across all samples. All subsequent analyses used the quality-control-filtered datasets.

### Preprocessing and Clustering

All preprocessing and clustering were performed in Python using Scanpy^46^. A Pearson-residuals framework that accounts for sequencing depth was used to identify highly variable genes and normalize expression values. Principal component analysis was then applied to the highly variable feature set, and sample-level batch effects were mitigated by integrating the principal components with harmonypy. The harmonypy-corrected embeddings were used to construct a k-nearest-neighbor graph, followed by UMAP for visualization. Clustering was performed with the Leiden algorithm to define transcriptionally coherent clusters.

### Cell-type annotation

Clustering of the integrated dataset resolved 40 clusters. We annotated clusters based on differentially expressed genes and canonical cell-type markers (Supplementary Tables 2–9). Epithelial cells, fibroblasts, macrophages, endothelial cells, and pericytes/smooth-muscle cells were further examined by subclustering. As several well-established markers for epithelial, fibroblast, and macrophage lineages are not included in the Xenium 5k gene panel, annotations for these compartments also incorporated results from imputation (detailed in a separate section), which infers the expression of panel-missing marker genes from public single-cell RNA-seq references.

### Differentially expressed genes analysis

For each analysis, raw counts were normalized to a fixed library size (10,000 counts per cell), followed by log transformation. Differentially expressed genes were then ranked between groups using the Wilcoxon rank-sum test, and the top-ranked genes per group were retained. For each candidate gene, we additionally computed the fraction of cells expressing the gene within each group (percentage of cells with non-zero expression) to aid interpretation. Differential expression results (gene names, scores, P values, multiple-testing adjusted P values, log fold changes, and detection fractions) were exported as tables (Supplementary Tables 2, 4–9).

### Imputation

Imputation refers to predicting the expression of genes that were not directly measured by leveraging gene–gene covariance learned from reference single-cell RNA-seq datasets. In practice, sample profiles from the Xenium 5k gene panel are mapped to a reference latent space, and gene-level information is transferred to infer panel-missing marker expression while preserving each cell’s identity and spatial context. As imputation inherently relies on model assumptions and reference data, it can be susceptible to library- and reference-specific biases. To mitigate these effects, we applied two independent imputation frameworks—Tangram^47^ and stAI^48^—together with two public single-cell RNA-seq references (GEO: GSE135893^7^ and GSE136831^14^), using only samples from idiopathic pulmonary fibrosis and control lungs. Predicted expression of panel-missing marker genes from these combinations was used solely to support cell-type annotation and not for differential testing or quantitative comparisons.

### Visualization of spatial cell-type annotations

We visualized cell-type annotations by rendering the cell-boundary polygons stored in the SpatialData shapes layer (“cell_boundaries”) with GeoPandas/Matplotlib. For each sample, polygons were loaded as a GeoDataFrame and assigned cell subtype labels by mapping polygon identifiers to the corresponding annotation table. Polygons were then grouped by subtype and plotted on a black background using a predefined color palette and subtype-specific hatching patterns.

### Neighborhood enrichment analysis across all cell subtypes

Neighborhood enrichment analysis was performed using the Squidpy. Briefly, a spatial neighbor graph was constructed from the two-dimensional cell coordinates using Delaunay triangulation. Squidpy was then used to quantify subtype–subtype neighborhood enrichment by comparing the observed frequency of neighboring subtype pairs to a permutation-based null model generated by repeatedly shuffling subtype labels. Enrichment was summarized as z-scores. For visualization, the resulting subtype-by-subtype z-score matrix was ordered by hierarchical clustering based on correlation distance with average linkage and displayed as a heatmap.

### Epithelial-centered neighborhood profiling

To quantify the local cellular milieu surrounding epithelial cells, we performed epithelial-centered neighborhood profiling. For each epithelial cell, we defined its neighborhood as all cells located within an 80-µm radius, based on cell-centroid distances in the 2D tissue plane. We counted neighboring cells by annotated subtype, yielding an epithelial cell-by-subtype neighborhood count matrix. Neighborhood feature matrices were processed using standard Scanpy workflows, including per-neighborhood total-count normalization, log transformation, dimensionality reduction, and graph-based clustering. Low-information neighborhoods were filtered before downstream analyses. Initial clustering identified 10 neighborhood clusters, which were subsequently grouped into seven broader neighborhood profiles according to shared patterns in the central epithelial cell identity, neighboring cell-type composition, and spatial-histologic distribution across tissue sections. The resulting profiles were annotated as near-normal alveolar, pre-fibrotic, fibrotic, metaplastic, bronchiolar, alveolar duct, and epithelial-rich.

### Inference of the pseudo-temporal continuum

Pseudo-temporal continuum inference was performed from epithelial-centered neighborhood profiles using the scFates^26^ package. Under a non-branching trajectory assumption consistent with putative unidirectional epithelial remodeling. The near-normal alveolar profile was designated as the root state, from which pseudotime was inferred. Pseudotime values were subsequently rescaled to a 0–1 range and defined as the continuum position. Changes in neighborhood cell-type counts along the continuum position were modeled within scFates, and enrichment dynamics along the continuum were evaluated using the pseudotime enrichment analysis implemented in decoupler^29^.

### Extension of continuum positions to nearby non-epithelial cells

To evaluate whether enrichment analysis based on the epithelial continuum could be affected by transcript spillover from adjacent cells, we extended the continuum framework to nearby non-epithelial cells. For each non-epithelial cell located within a 30-µm radius of epithelial cells, a continuum position was assigned on the basis of the continuum positions of neighboring epithelial cells using a distance-weighted scheme, such that closer epithelial cells contributed more strongly. This enabled continuum-based enrichment analysis to be performed across surrounding non-epithelial populations in addition to epithelial cells using decoupler. Comparing enrichment patterns between epithelial and non-epithelial cell types along the same continuum axis was then used to evaluate the potential contribution of transcript spillover to epithelial enrichment signals.

### Epithelial subtype-resolved pathway analysis along the pseudo-temporal continuum

For epithelial subtype-resolved analysis, the continuum was divided into five equal bins, and mean pathway activity scores were calculated for each epithelial subtype within each bin. Subtype-bin combinations representing <3% of epithelial cells in a given bin were treated as insufficiently represented and excluded from visualization, because estimates based on very small cell numbers are more susceptible to sampling noise and disproportionate influence from a few cells.

### Spatial distance-weighted ligand-receptor interaction analysis

To quantify epithelial cell-centered cell–cell communication, we calculated spatial distance-weighted ligand-receptor interaction scores using curated ligand–receptor pairs from the CellChat^30^ database, restricted to Secreted Signaling interactions. Only interactions for which all ligand and receptor genes were represented in the Xenium panel were retained. Per-cell expression values were library-size normalized, and ligand and receptor activities were summarized at the single-cell level using the geometric mean of the corresponding genes. Spatial neighborhoods were defined within a 300-µm radius around each epithelial cell, and neighboring cells were weighted according to their distance from the epithelial cell using a Gaussian kernel. Directional interaction scores were then computed for both neighboring-cell-to-epithelial and epithelial-to-neighbor signaling. These scores were subsequently used in three downstream analyses: aggregation at the epithelial subtype level to characterize subtype-resolved interaction patterns, stratification of epithelial cells into signal-positive and signal-negative groups based on whether the corresponding interaction score was greater than zero or equal to zero, and aggregation across epithelial continuum-position bins to examine dynamic changes in cell–cell communication along the fibrotic continuum. Detailed mathematical definitions and analytical procedures are provided in Supplementary Note.

### Analysis of transcriptional programs associated with inferred interaction signals

To assess transcriptional programs associated with inferred incoming signals, we performed a binary comparison for each interaction of interest, including TGFβ, HGF–MET, PERIOSTIN, and C5–C5AR1. For each interaction, epithelial cells were assigned to a signal-positive group if the inferred incoming interaction score was greater than 0 and to a signal-negative group if the score was 0. To prevent downstream enrichment analysis from being driven by the ligand- and receptor-encoding genes used to define the interaction scores, these genes were excluded from the 5,001-gene Xenium panel before pseudo-bulk aggregation. Counts were then aggregated into pseudobulk profiles using decoupler by summing expression values within each sample, epithelial subtype, and signal group. Pseudobulk samples containing fewer than 10 cells or fewer than 1,000 total counts were excluded before downstream analysis. Differential expression analysis was performed using PyDESeq2^49^ either across all epithelial cells or separately within each epithelial subtype. In both analyses, sample was included as a covariate together with signal status, and Wald statistics for the contrast between signal-positive and signal-negative groups were obtained using PyDESeq2. Genes were subsequently ranked according to the resulting Wald statistics and analyzed with GSEApy^50^ using the Hallmark gene sets. For TGFβ and HGF–MET interactions, enrichment of the EPITHELIAL_MESENCHYMAL_TRANSITION term was specifically examined and visualized using the normalized enrichment score (NES) and nominal P value. For the remaining interactions, all Hallmark terms were tested, and the results were summarized using NES and false discovery rate (FDR) q values.

### Immunofluorescence staining

Formalin-fixed paraffin-embedded (FFPE) lung tissue sections from patients with IPF were incubated at 60°C for 30 min and deparaffinized. Sections were subjected to antigen retrieval in citrate buffer at 121℃ for 15 min and cooled to room temperature. After three washes in PBS, tissue areas were circled with an A-PAP PEN (#APAP-M, Daido Sangyo) and blocked with Blocking One Histo (#06349-64, Nacalai tesque) for 10 min. Tissue sections were washed once in 0.1% PBS-T and incubated overnight at 4℃ in a humidified chamber with the following primary antibodies: rabbit anti-periostin (1:200; #ab14041, Abcam), rabbit anti-pan-cytokeratin (1:200; #bs-2190R, Bioss), mouse anti-α-SMA (1:200; #A2547, Sigma), and mouse anti-integrin αVβ3 (1:100; #MAB1976, Merck). The next day, sections were washed three times in 0.1% PBS-T and incubated with fluorescently labeled secondary antibodies (1:500, #A21206 and #A31573, Invitrogen) and DAPI solution (1:500; #19178-91, Nacalai tesque) for 1 h in the dark at room temperature. After three additional washes, tissue sections were mounted (#DAI-DM-01, Daido Sangyo) and kept in the dark at 4°C until further analysis. Images were acquired using a BZ-X810 fluorescence microscope (Keyence).

## Code availability

Under preparation.

## Data availability

Under preparation.

## Author contributions

S.N., and K.T., contributed to the conception and design of the study. K.T., K.N., M.H., and Y.O. contributed to sample selection. M.H. and Y.O. performed pathological evaluation and tissue preparation. T. Takenaka contributed to the coordination of surgical specimen acquisition and preparation. Y.Y. performed immunofluorescence staining and generated the corresponding validation data. S.N. performed the data analysis. S.N., K.T., Y.Y., T. Takano, K.N., M.H., Y.O., and I.O.contributed to data interpretation. S.N. drafted the original manuscript. S.N., T. Takano, and K.T. reviewed and edited the manuscript. K.T. and I.O. organized and supervised the study.

