## Supplementary figures and images for "Trajectory inference of epithelial-centered neighborhood profiles reconstructs a pseudo-temporal continuum in idiopathic pulmonary fibrosis"

### Extended Data Figure

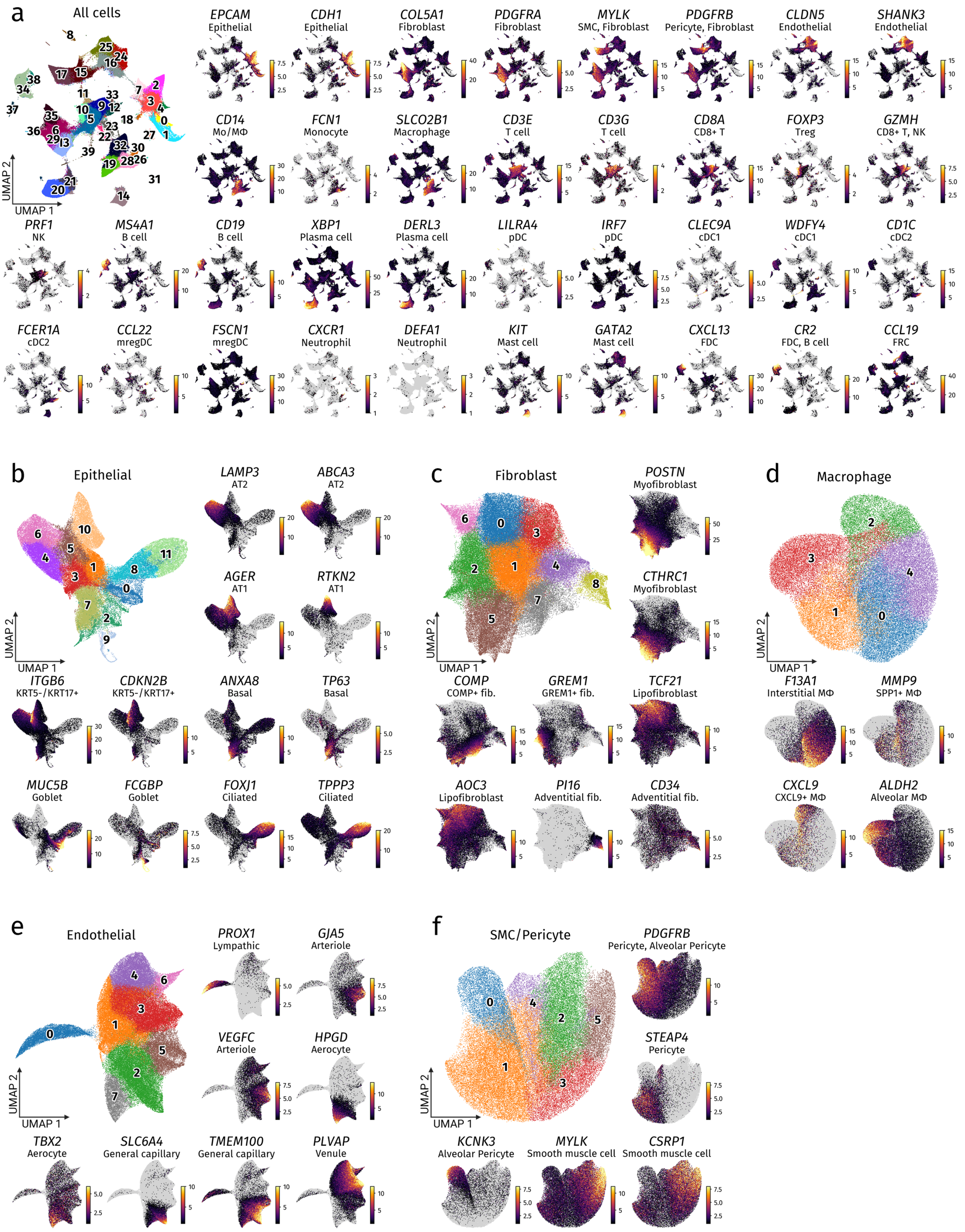

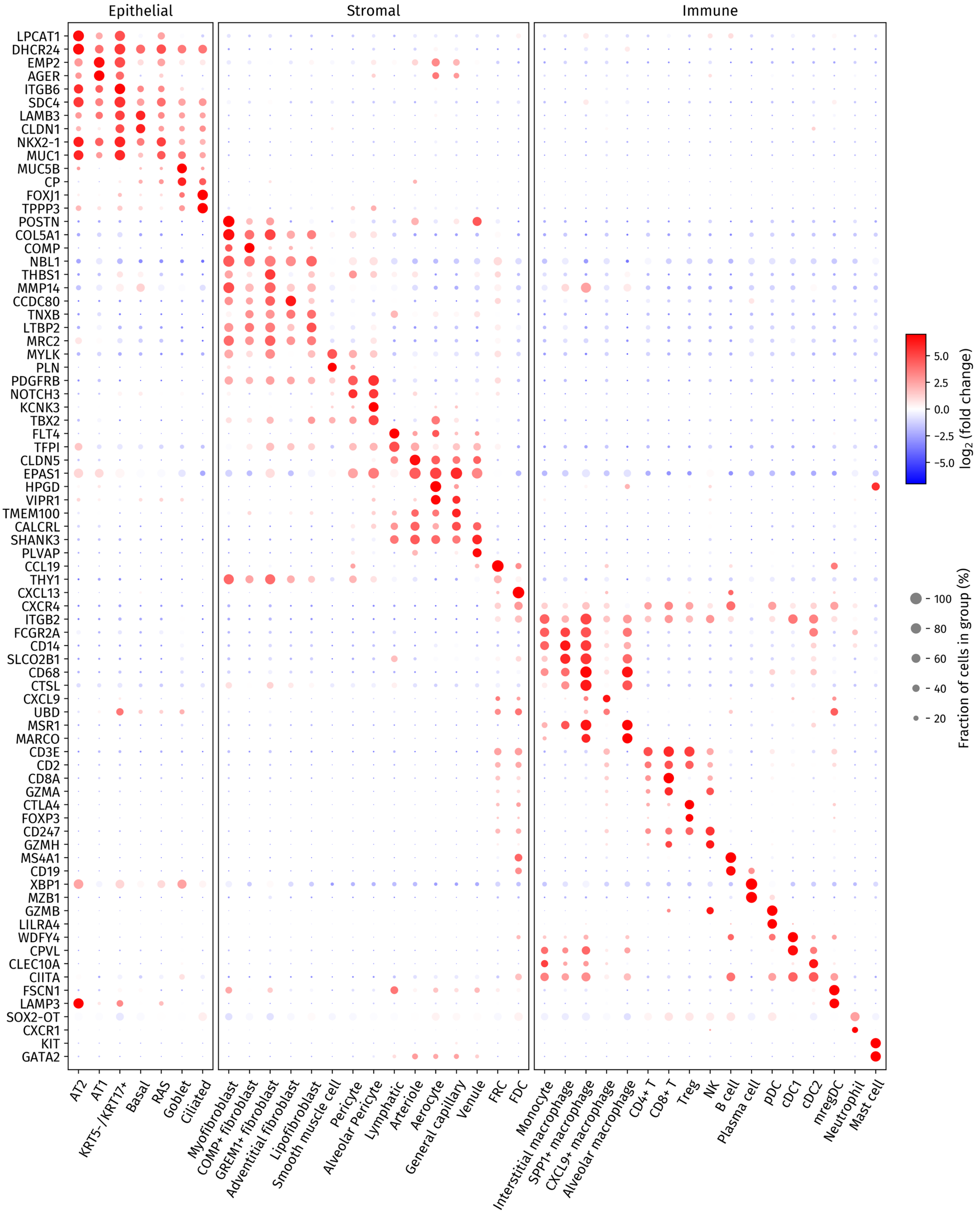

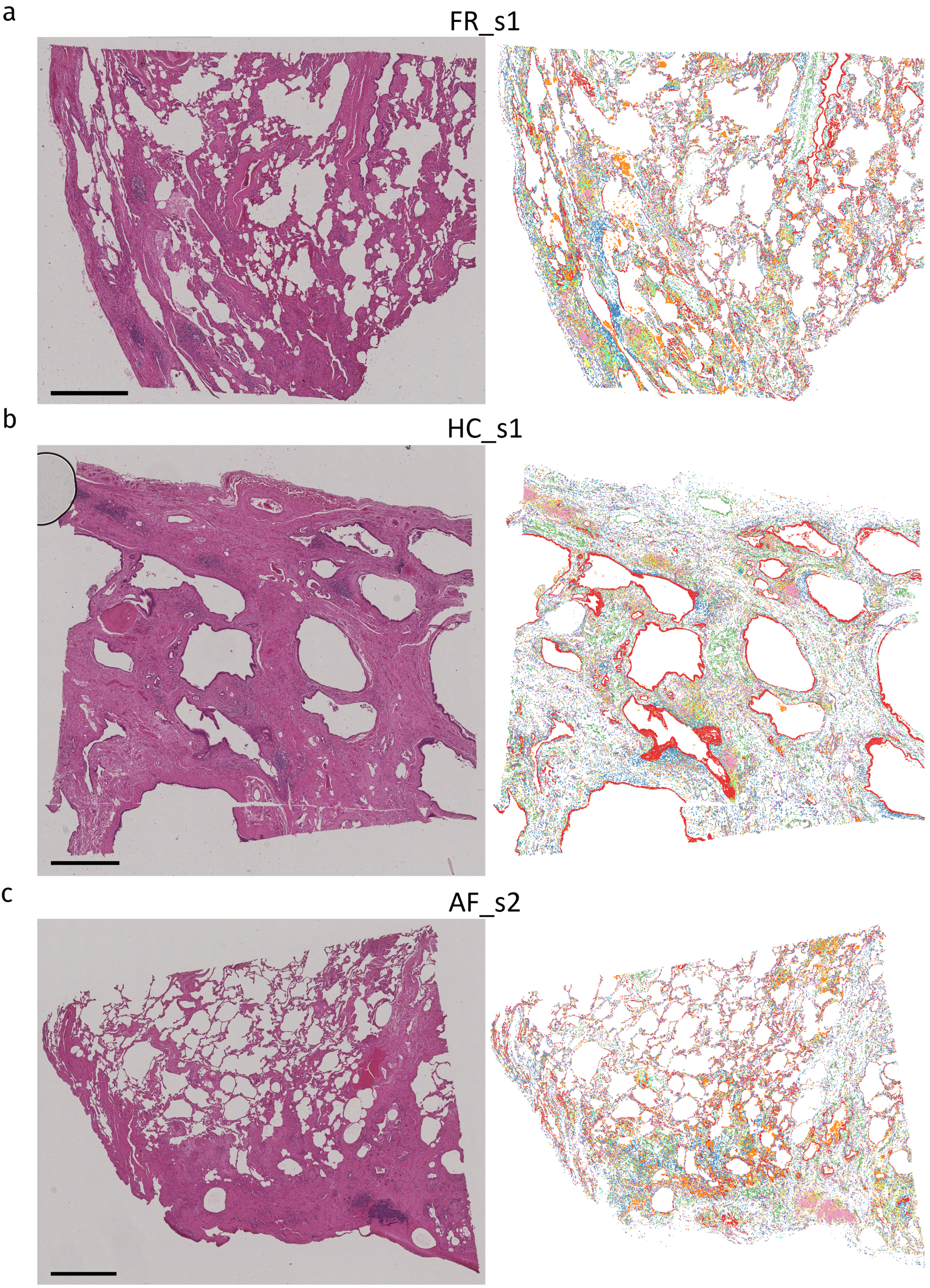

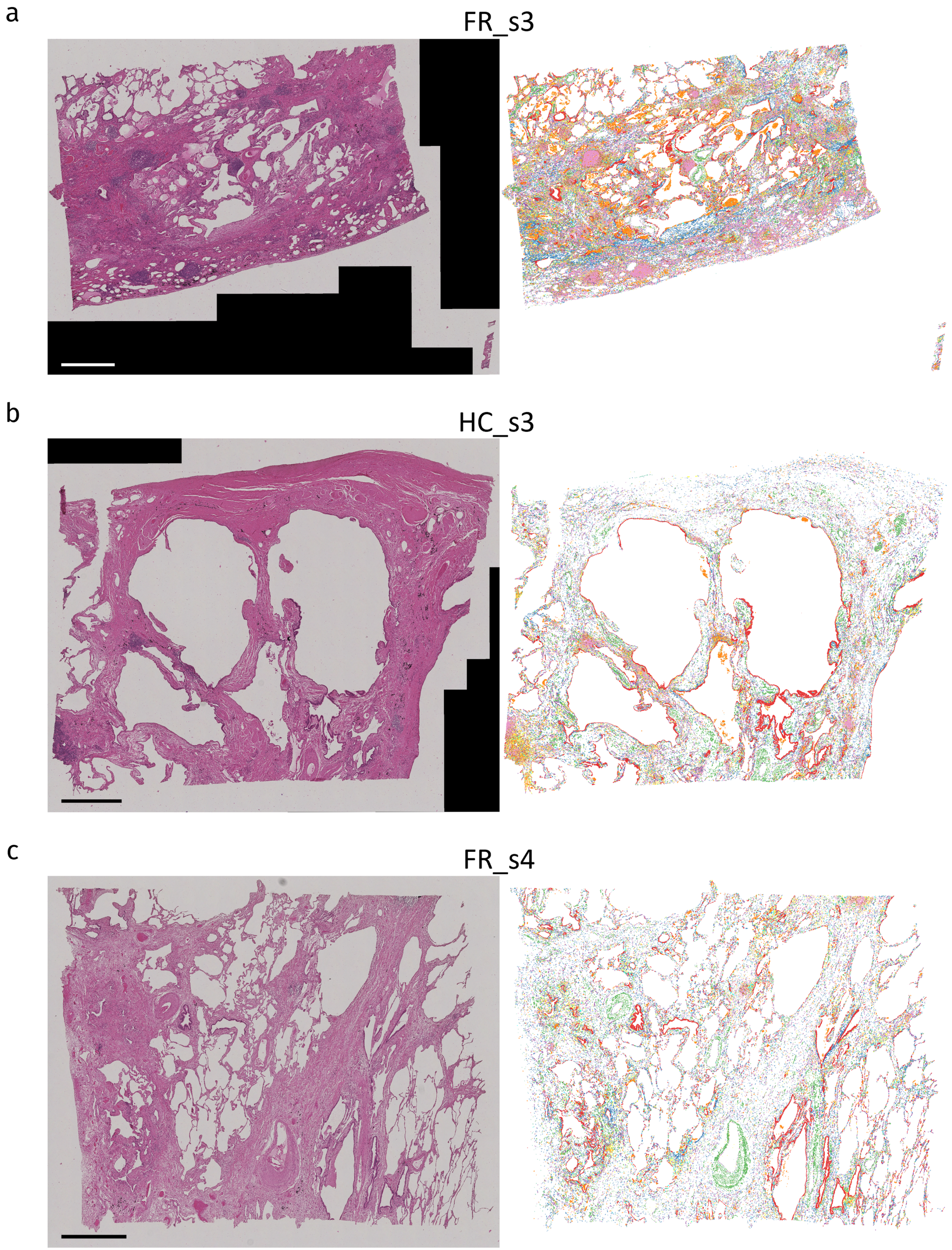

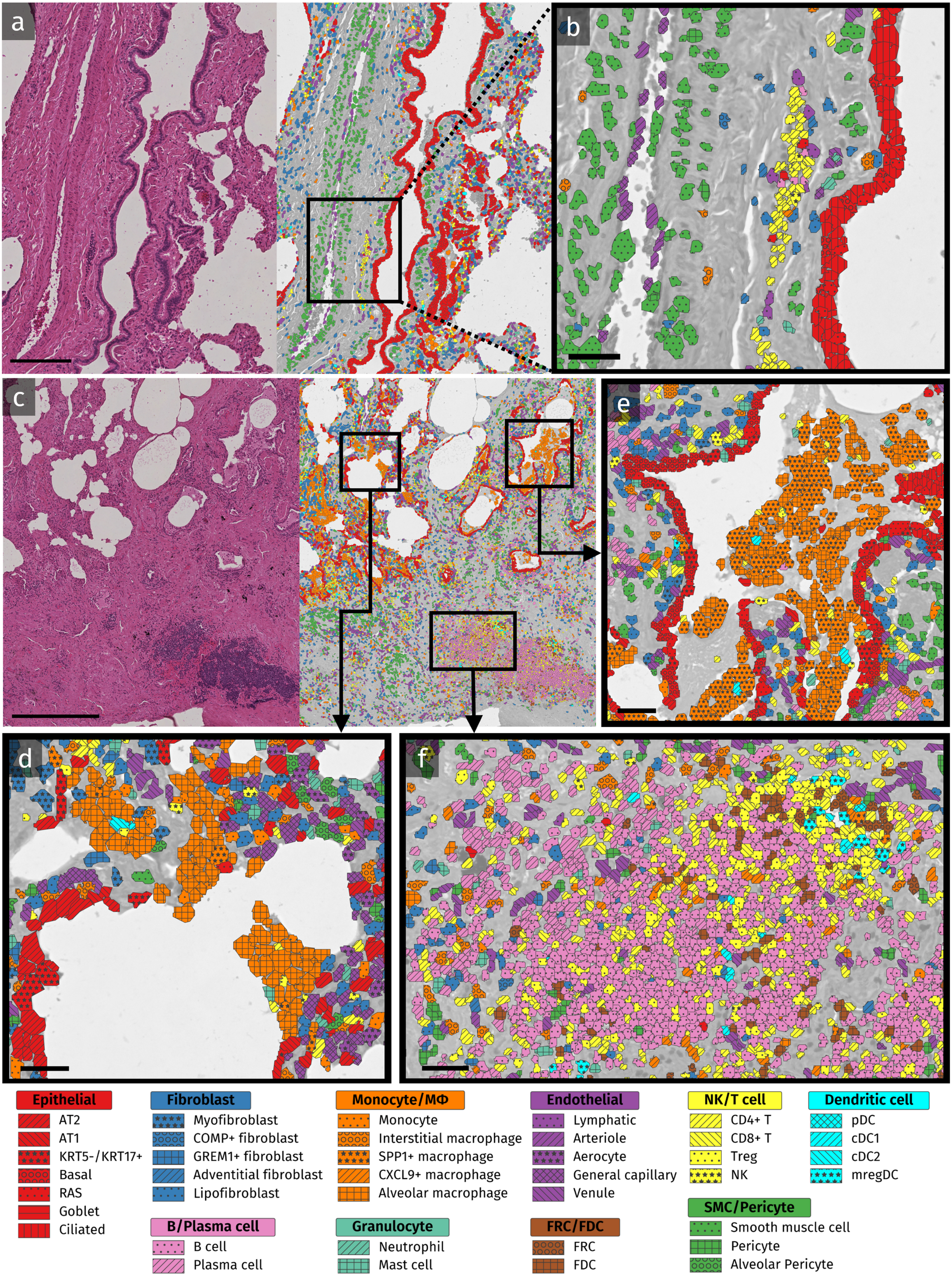

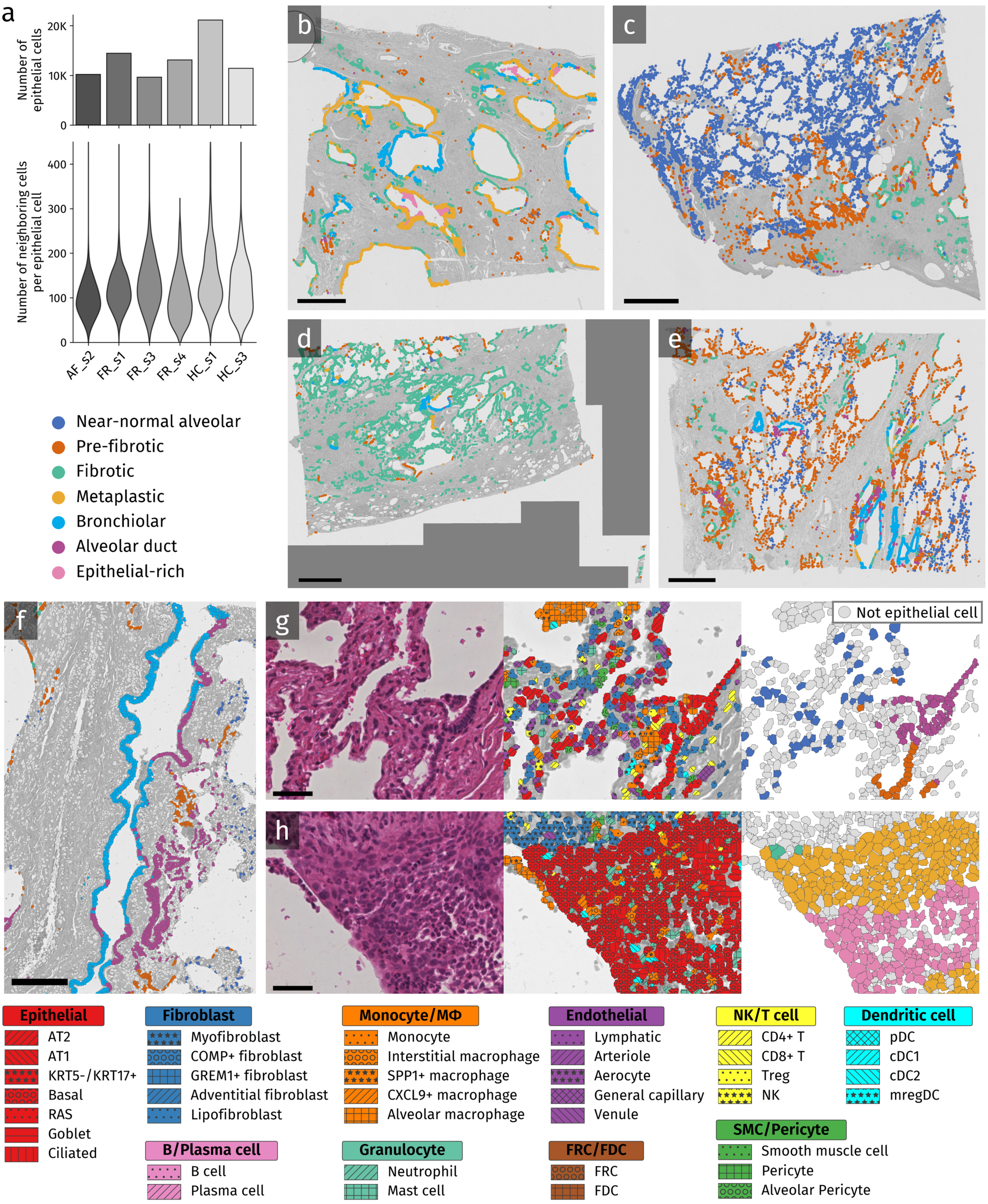

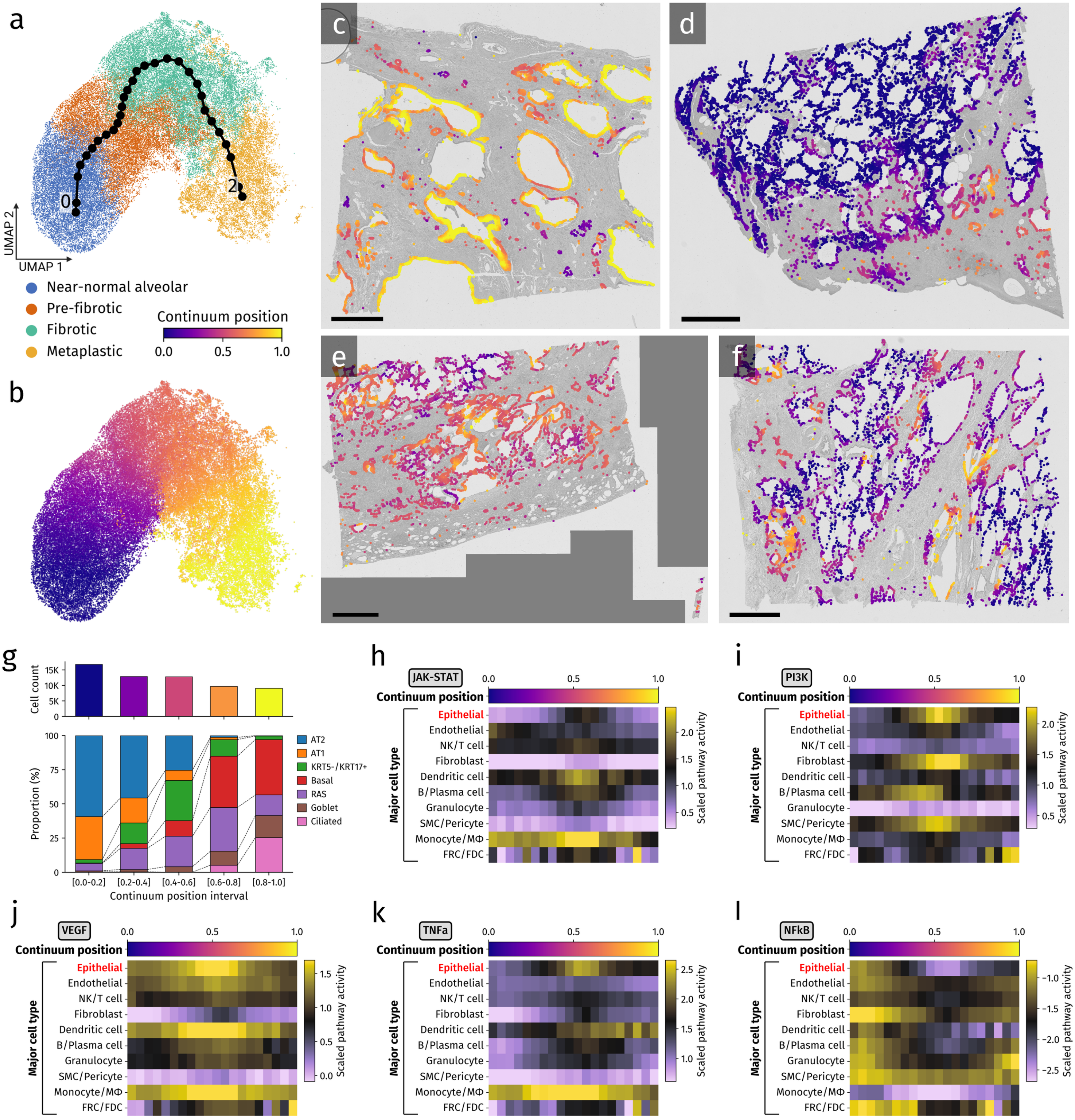

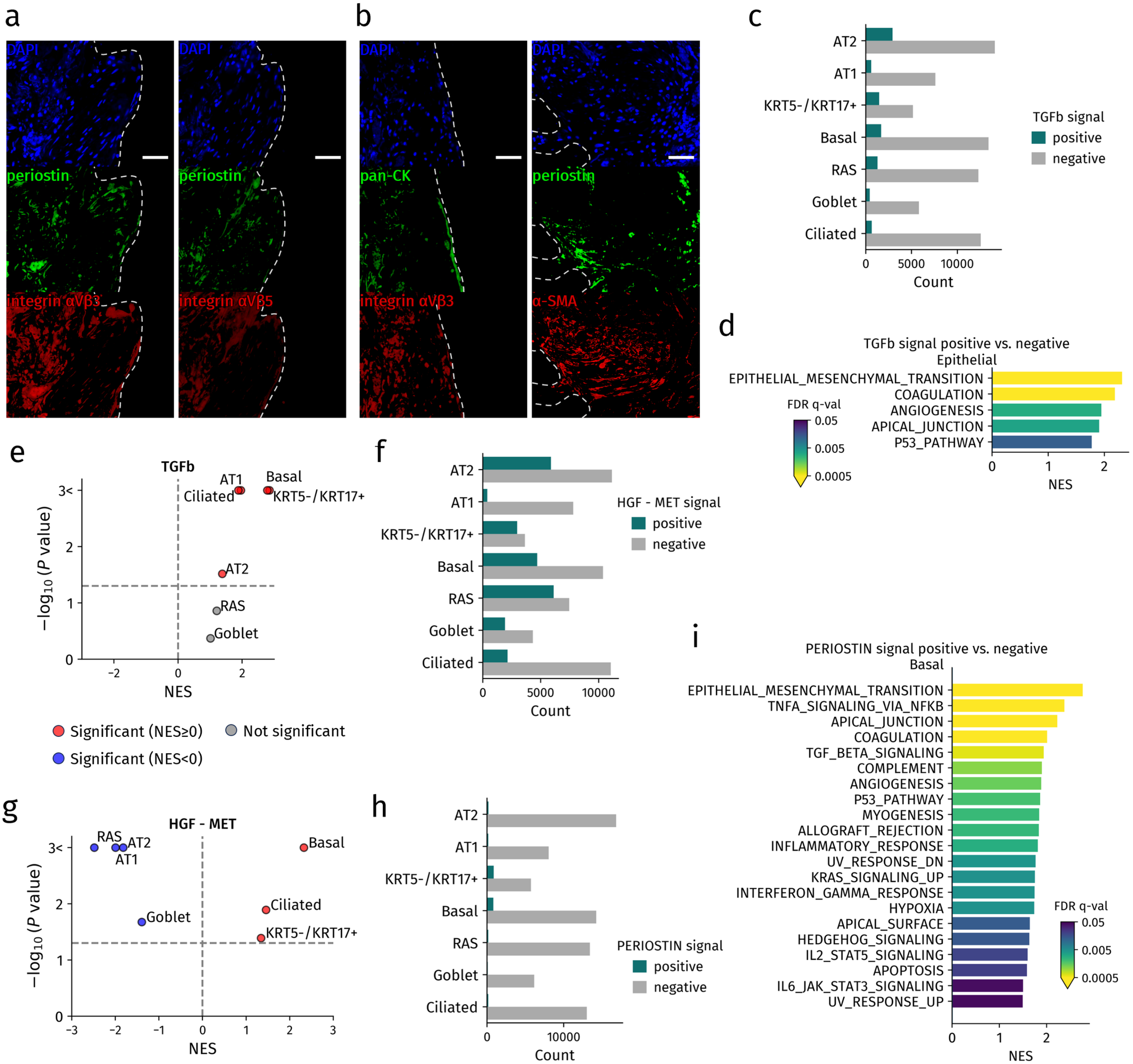

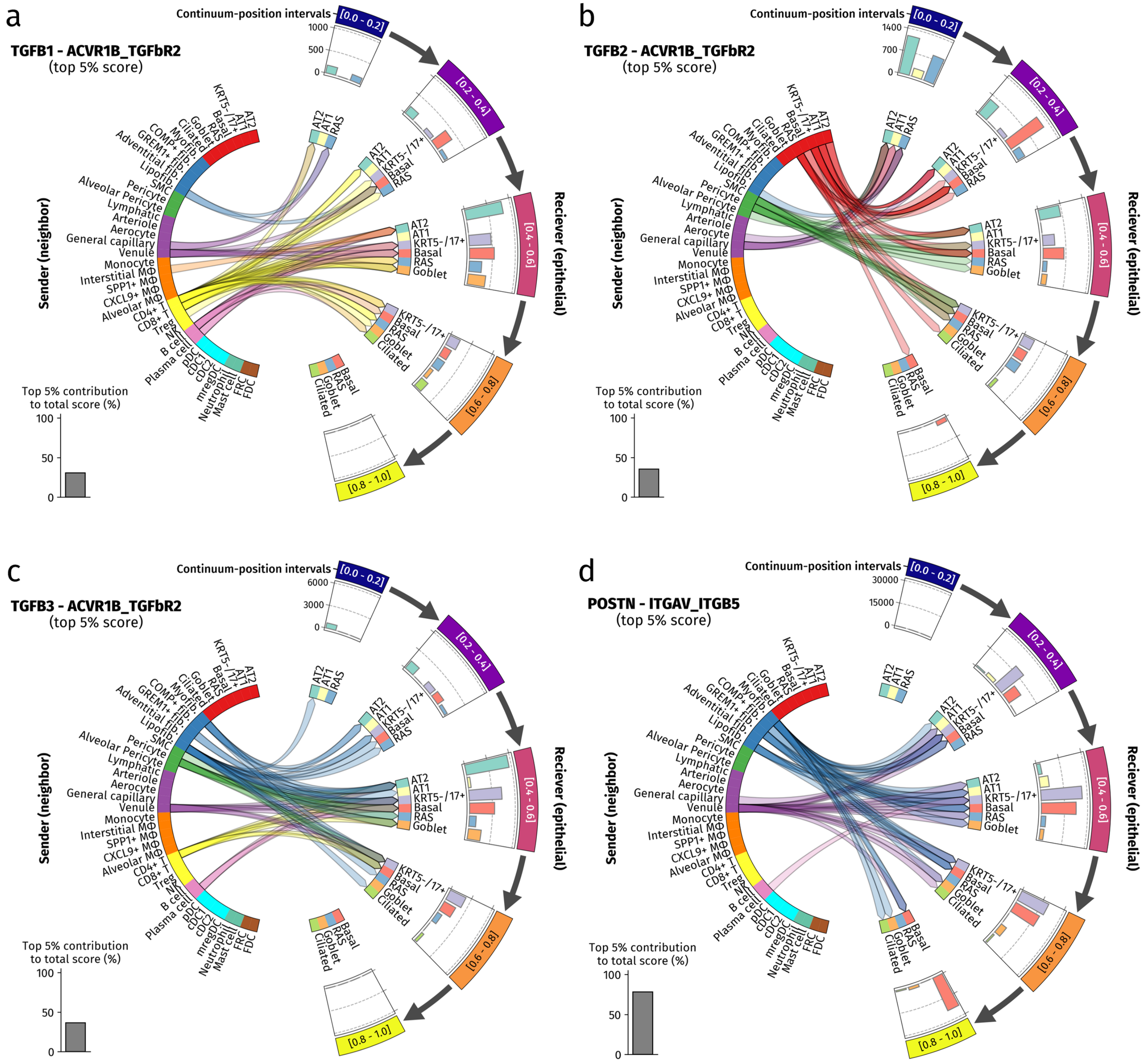

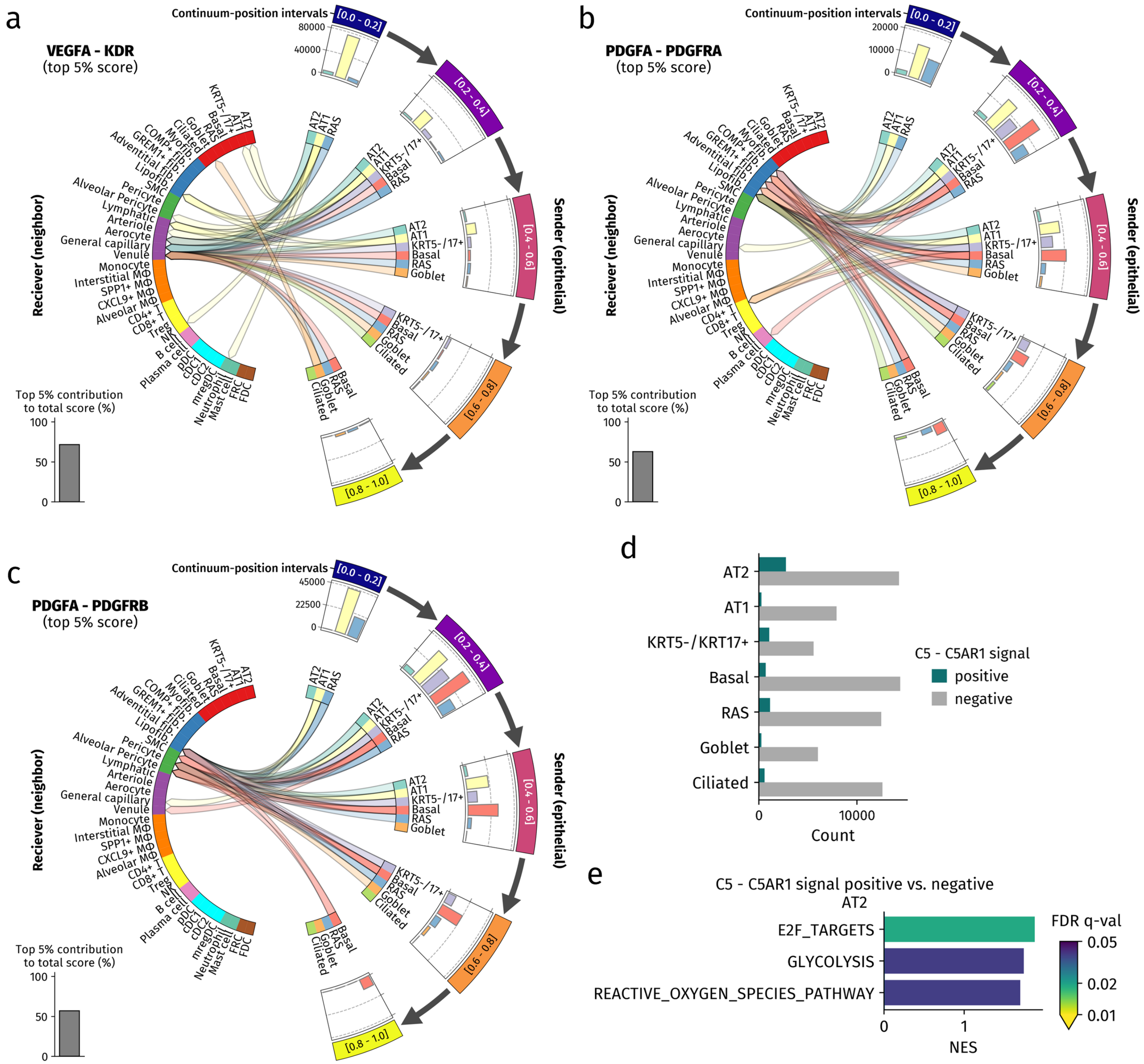

### Supplementary Figure

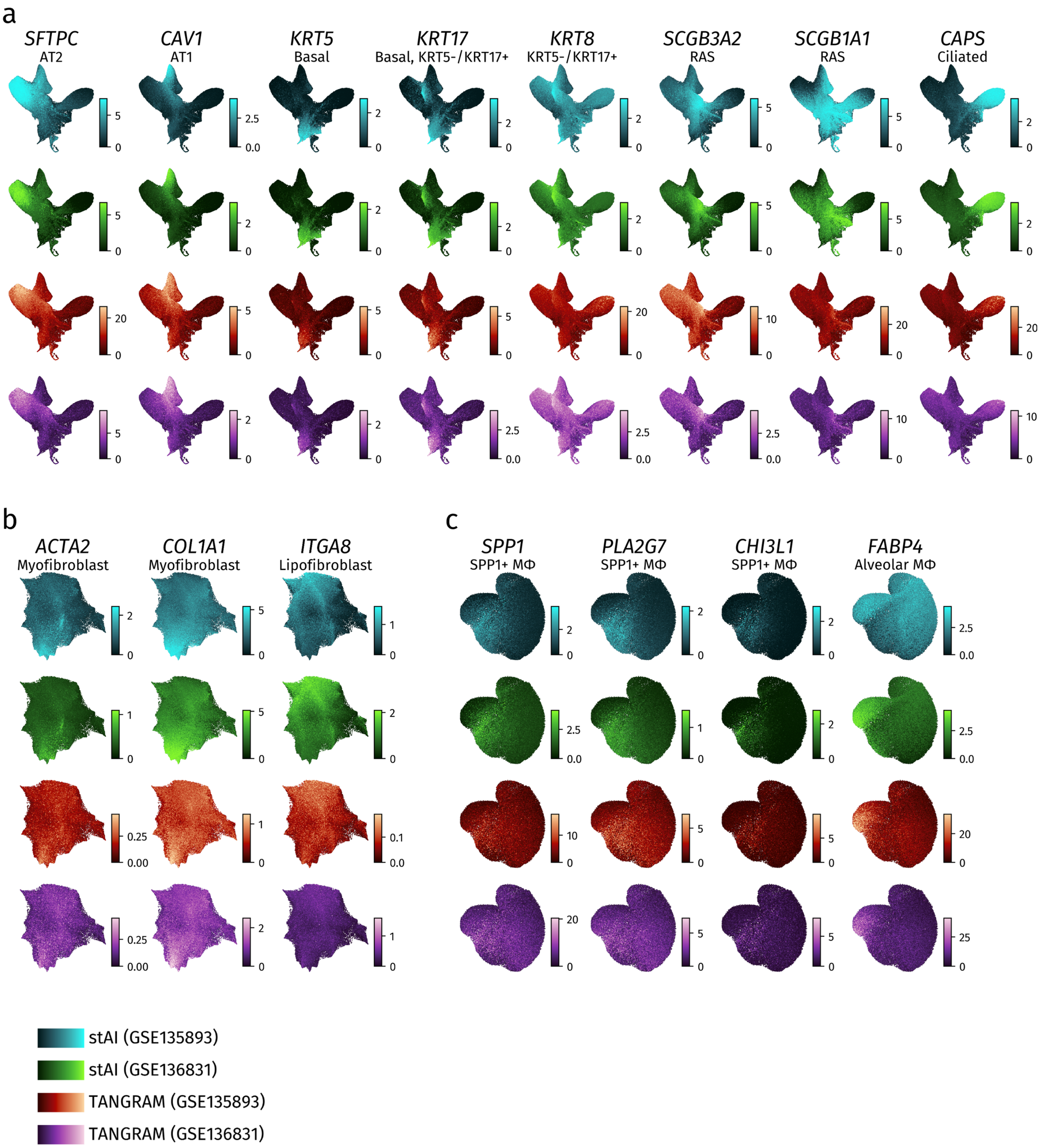

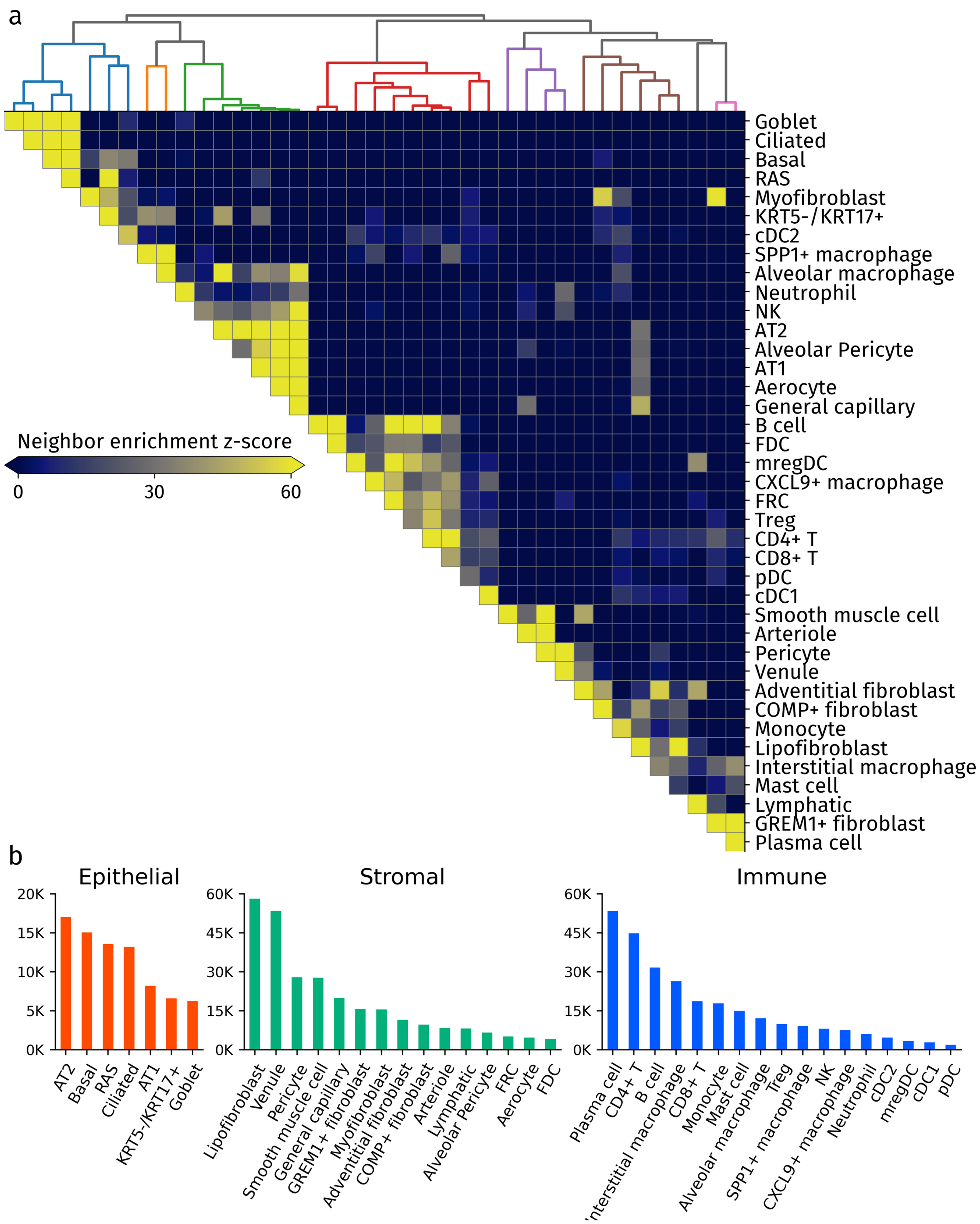

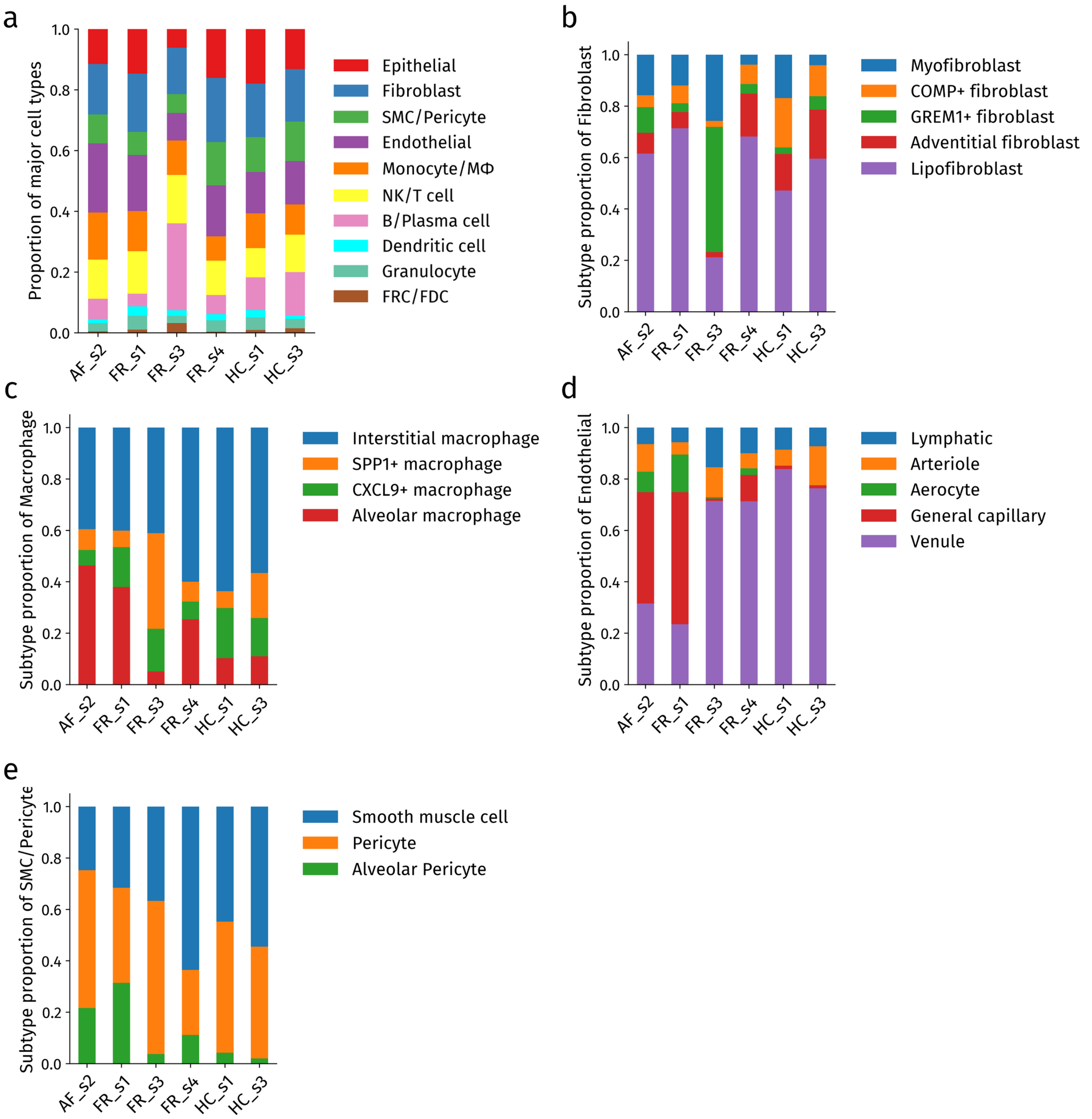

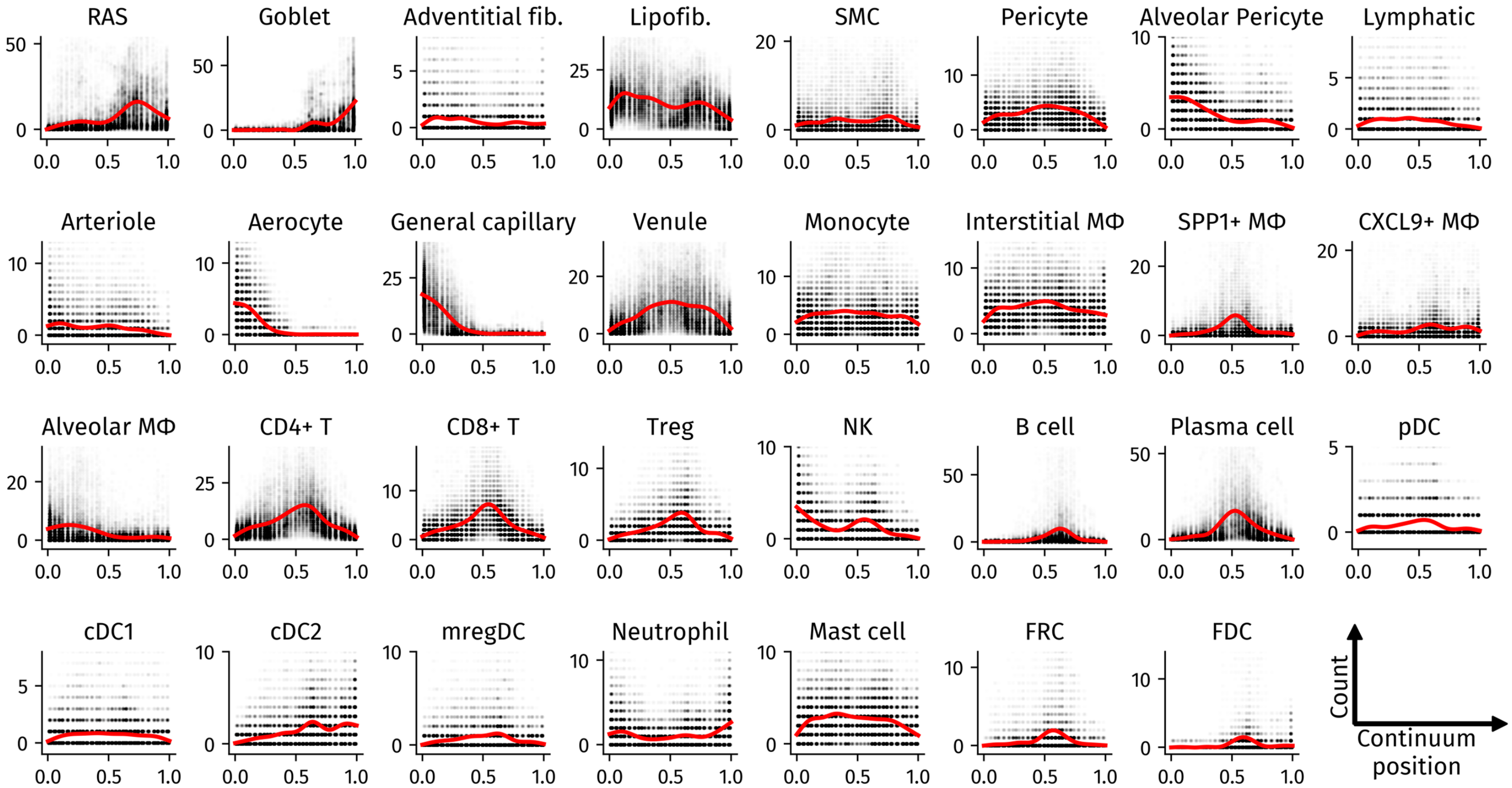
