## Supplementary Information for "Trajectory inference of epithelial-centered neighborhood profiles reconstructs a pseudo-temporal continuum in idiopathic pulmonary fibrosis"

### Extended Data Figure Legends

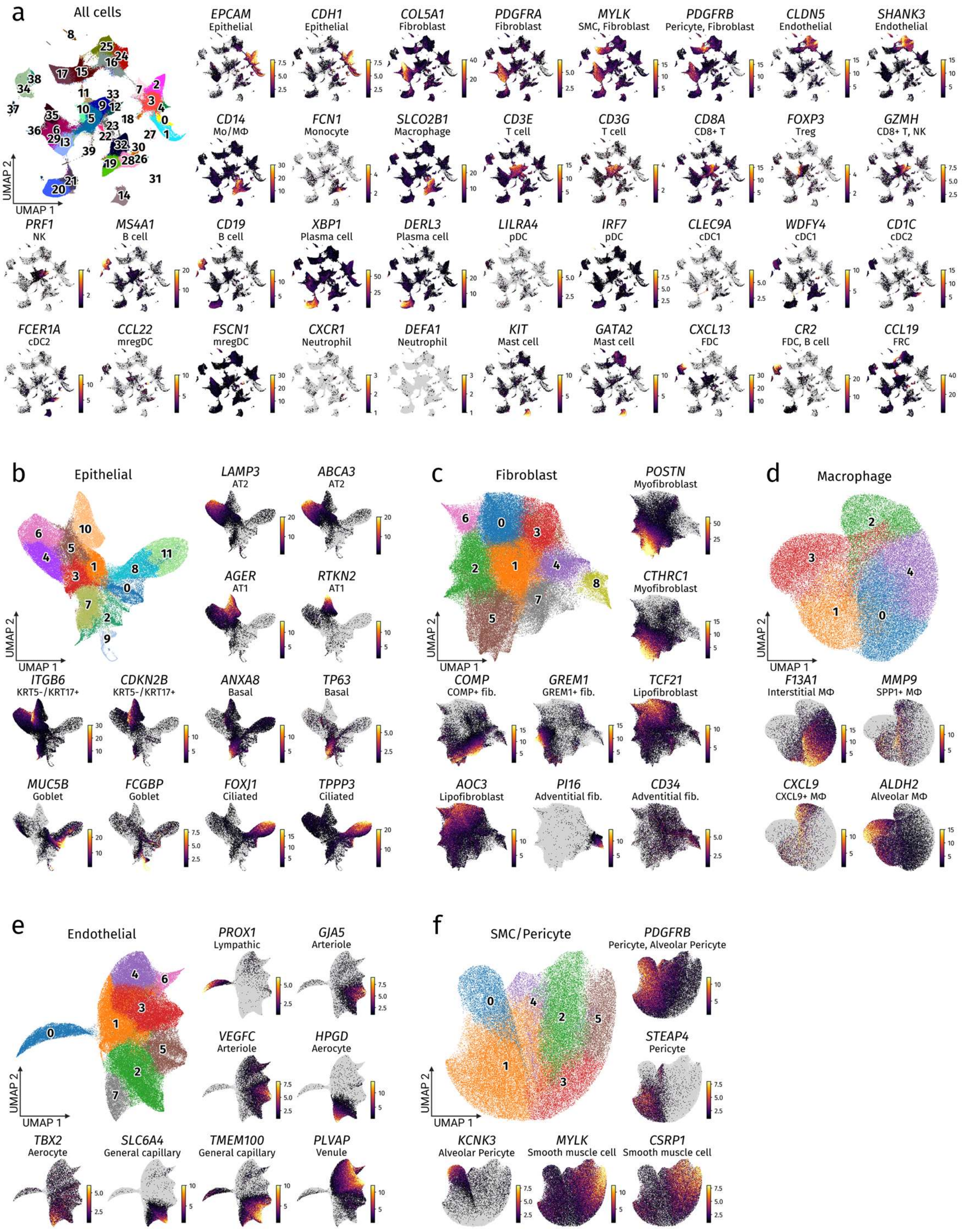


**Extended Data Fig. 1 | Marker gene expression supporting cel-type and subtype annotation. a,** UMAP embedding of all cells and feature plots of representative marker genes used for initial cell-type annotation. **b–f,** UMAP embeddings and feature plots of representative marker genes used for lineage-specific subtype annotation of epithelial cells (b), fibroblasts (c), macrophages (d), endothelial cells (e), and smooth muscle cells/pericytes (f). Color intensity indicates normalized gene expression. DC, dendritic cell; pDC, plasmacytoid dendritic cell; cDC1, conventional dendritic cell type 1; cDC2, conventional dendritic cell type 2; mregDC, mature dendritic cell enriched in immunoregulatory molecules; FRC, fibroblastic reticular cell; FDC, follicular dendritic cell; SMC, smooth muscle cell; RAS, respiratory airway secretory; Mo/MΦ, monocyte/macrophage.

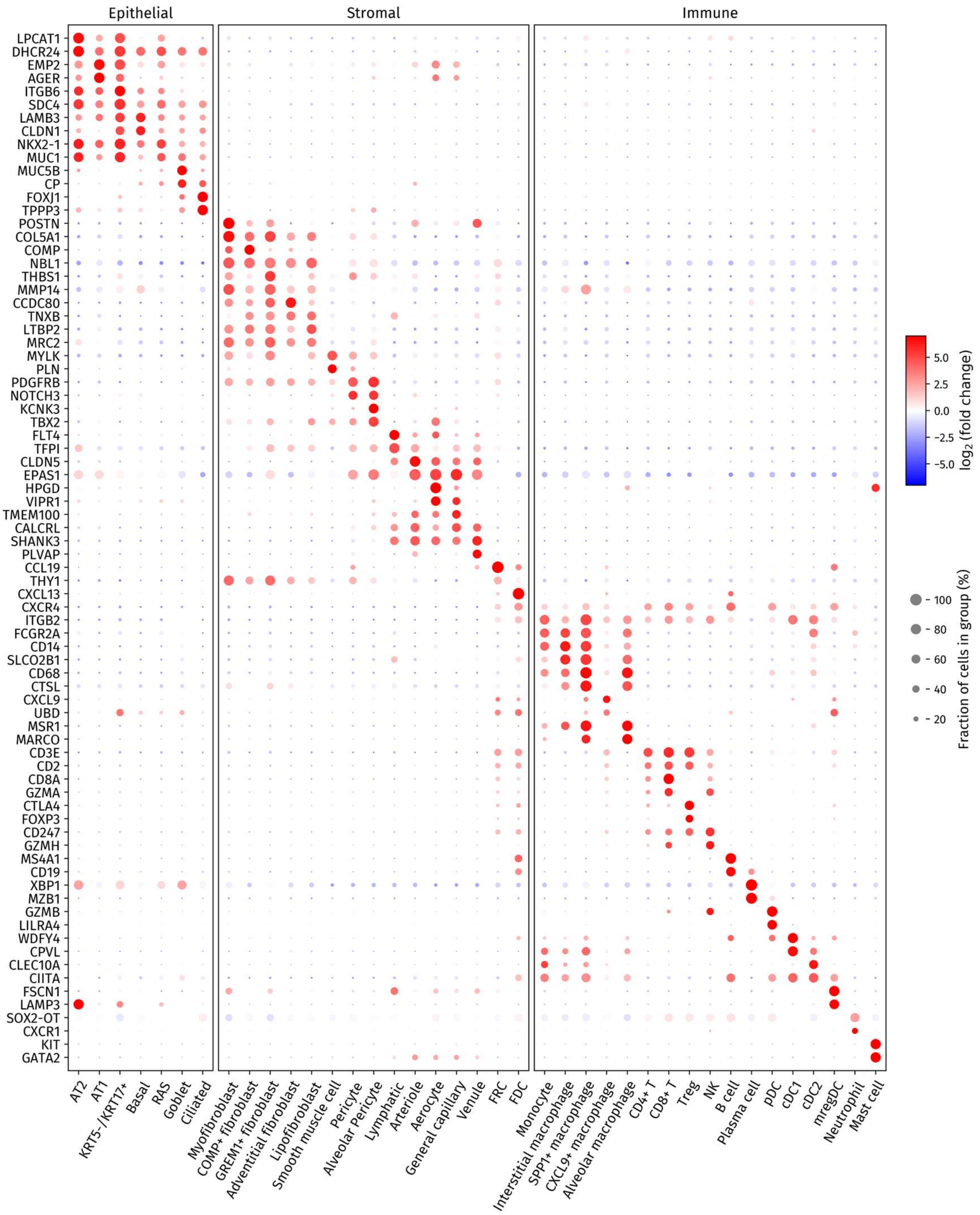


**Extended Data Fig. 2 | Differentially expressed marker genes across annotated cell subtypes.** Dot plot showing representative differentially expressed genes in annotated cell subtypes. For each subtype, the top two genes were selected while avoiding duplicate genes across subtypes. Columns indicate annotated subtypes grouped into epithelial, stromal, and immune compartments. Dot color indicates log2 fold change, and dot size indicates the fraction of cells expressing each gene within each subtype.

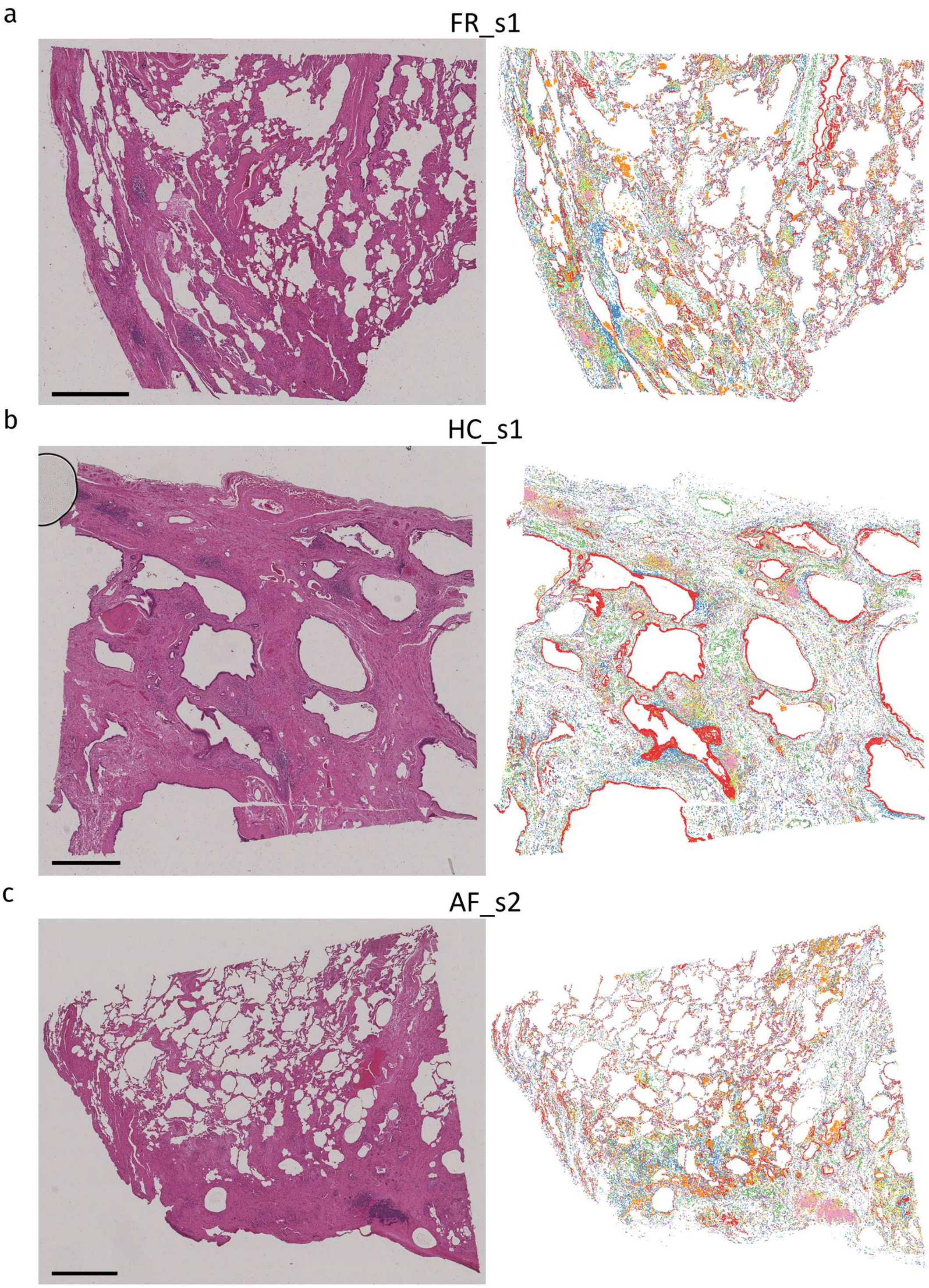


**Extended Data Fig. 3 | Spatial distribution of major cell types in FR_s1, HC_s1, and AF_s2. a–c,** H&E images and corresponding spatial maps of major cell types in FR_s1 (a), HC_s1 (b), and AF_s2 (c). Left panels show H&E-stained tissue sections, and right panels show Xenium/ProSeg-derived cell maps overlaid on grayscale tissue images. Cells are color-coded by major cell type using the same color scheme as in Fig. 1h–j. Scale bars, 1,000 µm.

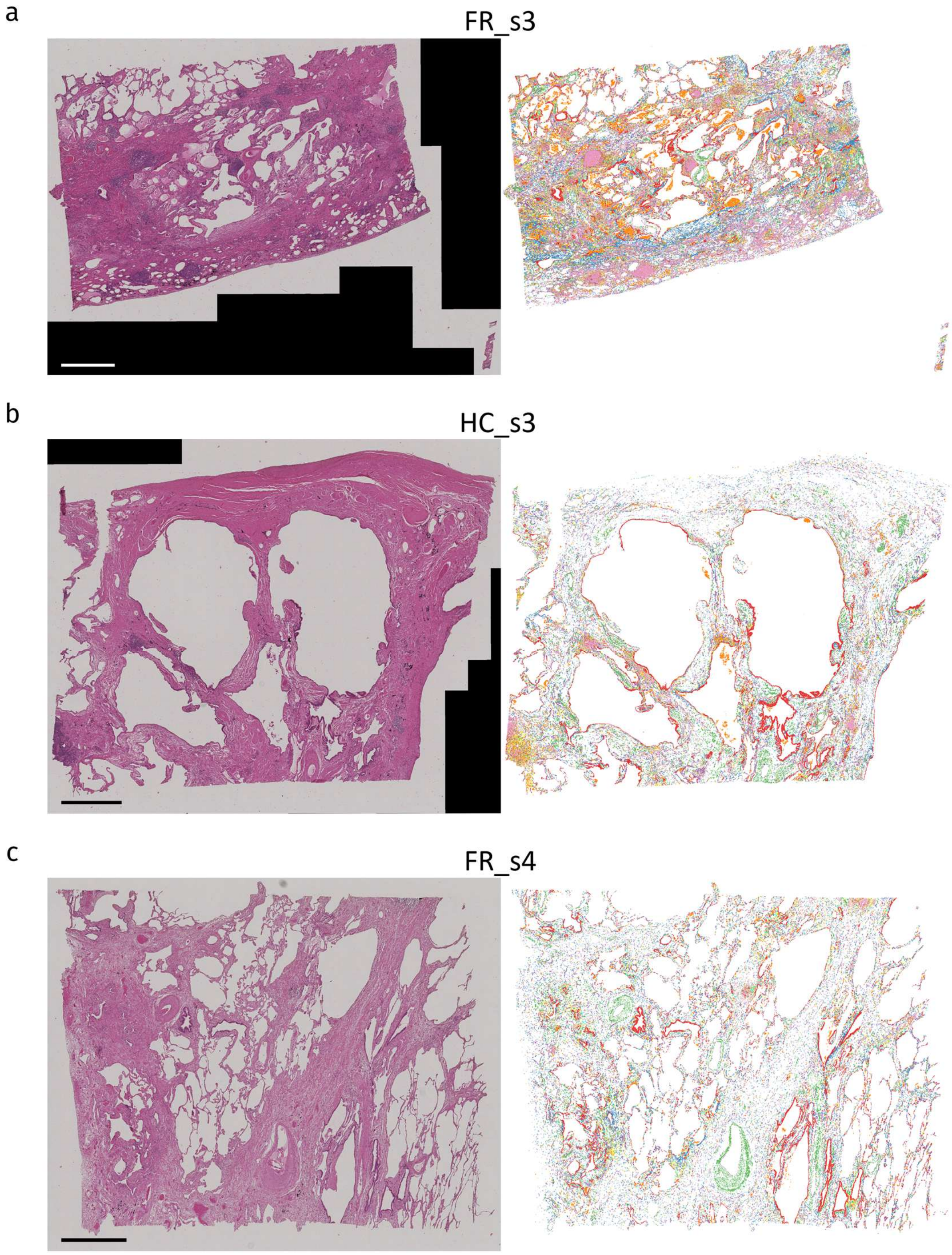


**Extended Data Fig. 4 | Spatial distribution of major cell types in FR_s3, HC_s3, and FR_s4. a–c,** H&E images and corresponding spatial maps of major cell types in FR_s3 (a), HC_s3 (b), and FR_s4 (c). Left panels show H&E-stained tissue sections, and right panels show Xenium/ProSeg-derived cell maps overlaid on grayscale tissue images. Cells are color-coded by major cell type using the same color scheme as in Fig. 1h–j. Scale bars, 1,000 μm.

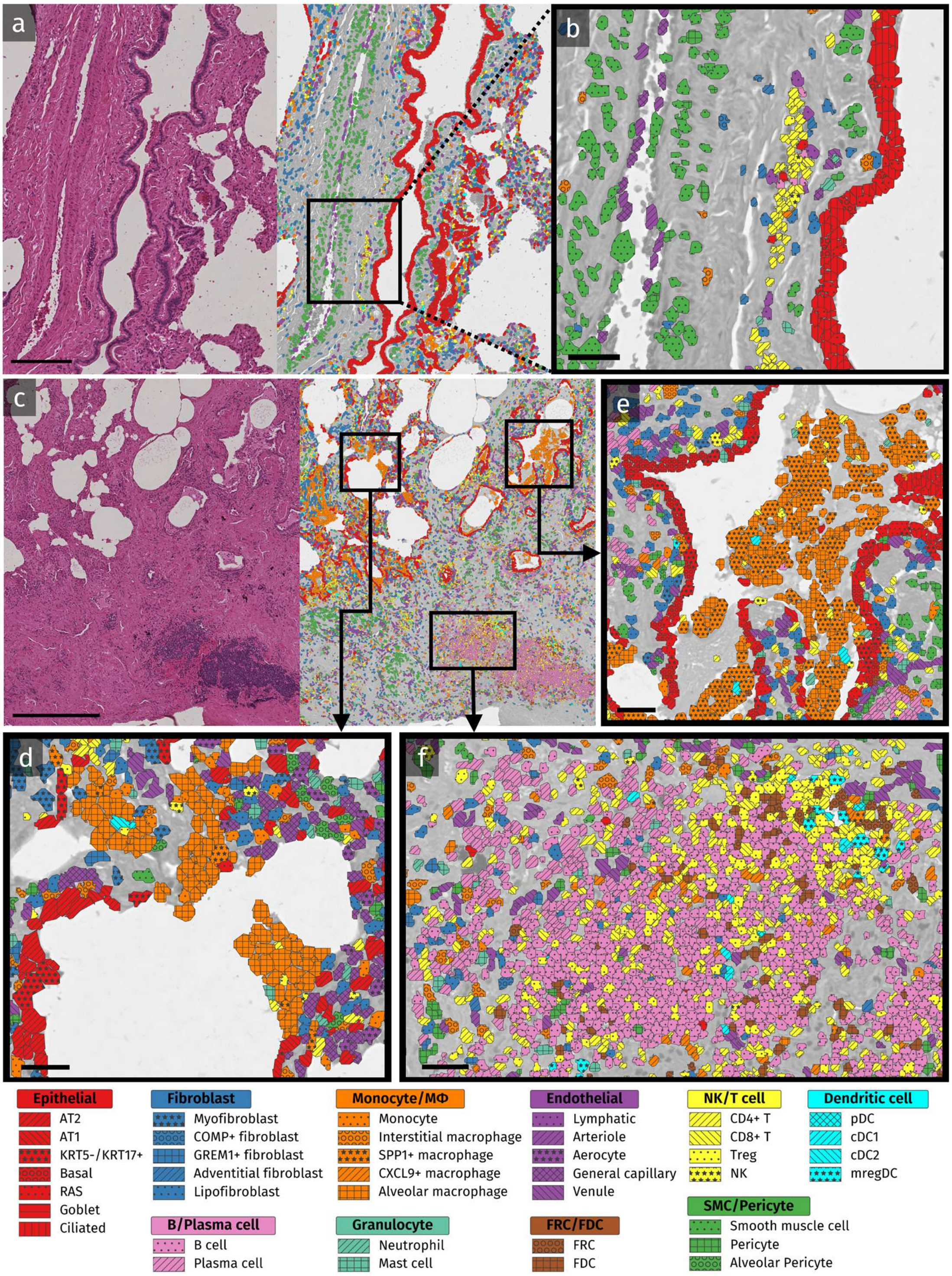


**Extended Data Fig. 5 | Spatial annotation of anatomical and immune structures in IPF lung tissues. a,** H&E image and corresponding spatial map of annotated cell types in FR_s1, showing a bronchus running alongside a bronchial artery. **b,** Magnified view of the boxed region in **a**, showing subtype-level annotation indicated by hatching patterns. The region includes airway epithelium, vascular structures, and lymphatic endothelial cells. **c,** H&E image and corresponding spatial map of a selected region of AF_s2. **d–f,** Magnified views of the boxed regions in **c**, showing alveolar macrophages within alveolar spaces (d) *SPP1*$^{+}$ macrophage accumulation within alveolar spaces (e), and a tertiary lymphoid structure within the lung interstitium (f). Cell colors indicate major cell types, and hatching patterns indicate annotated subtypes as shown in the legend. Scale bars, 200 µm (a), 500 µm (c), and 40 µm (b, d–f).

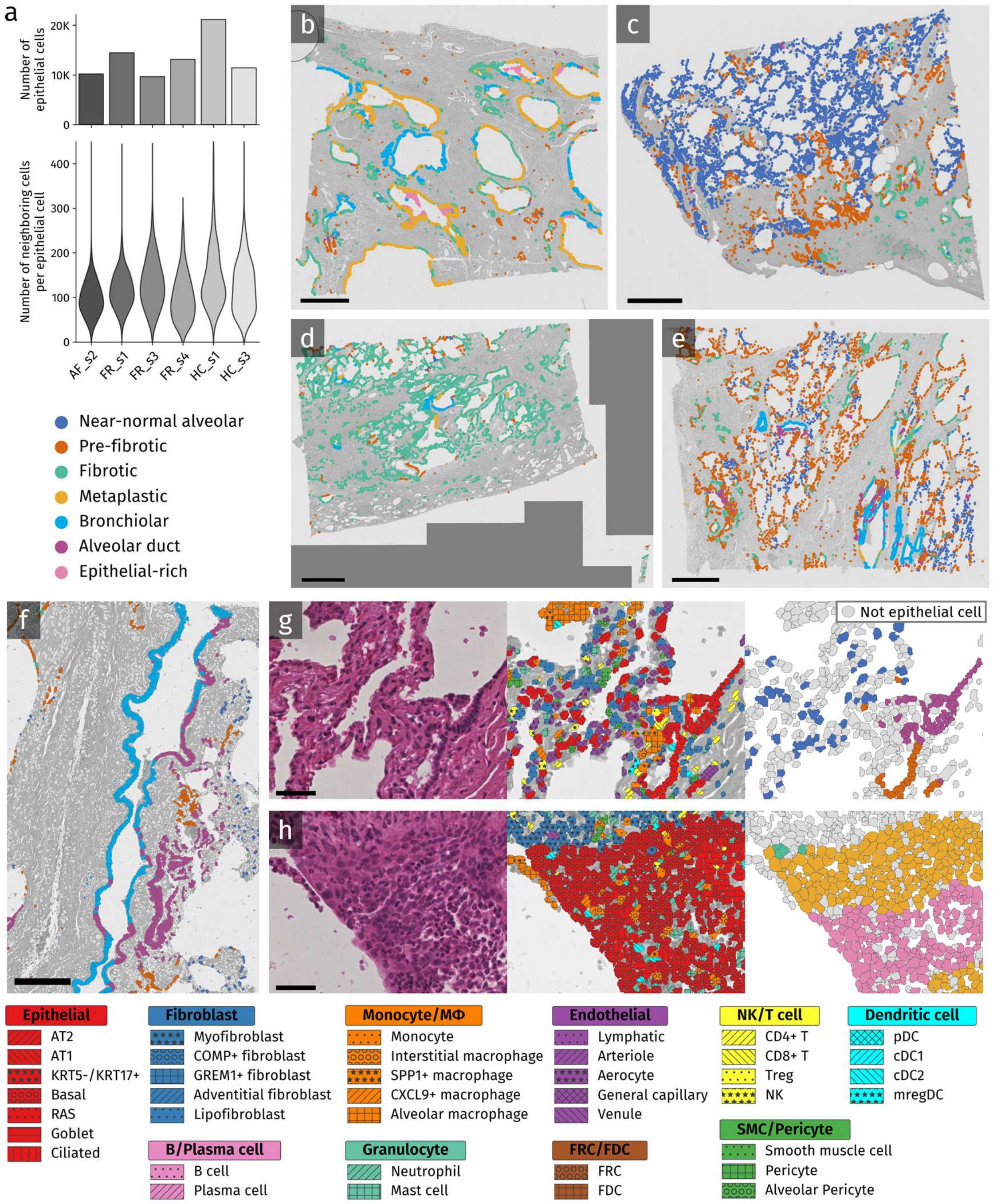


**Extended Data Fig. 6 | Additional features and spatial examples of epithelial-centered neighborhood classes. a,** Bar plots showing the number of epithelial cells in each sample (top) and violin plots showing the distribution of neighboring-cell counts within an 80-μm radius of each epithelial cell (bottom). **b–e,** Spatial maps of epithelial cells color-coded by neighborhood class overlaid on grayscale tissue images in HC_s1 (b), AF_s2 (c), FR_s3 (d), and FR_s4 (e). **f,** Spatial maps of epithelial cells color-coded by neighborhood class in a bronchial region corresponding to Extended Data Fig. 5a. **g, h,** Left panels show H&E images, middle panels show Xenium/ProSeg cell-type segmentation maps with colors and hatching indicating annotated cell subtypes, and right panels show neighborhood-class maps in which epithelial cells are color-coded by neighborhood class and non-epithelial cells are shown in gray. Panel (g) shows alveolar ducts adjacent to near-normal alveolar areas in FR_s1, and panel (h) shows luminal/apical side of markedly expanded airway epithelium in FR_s3. Scale bars, 1000μm (b–e), 200 μm (f), and 50 μm (g, h).

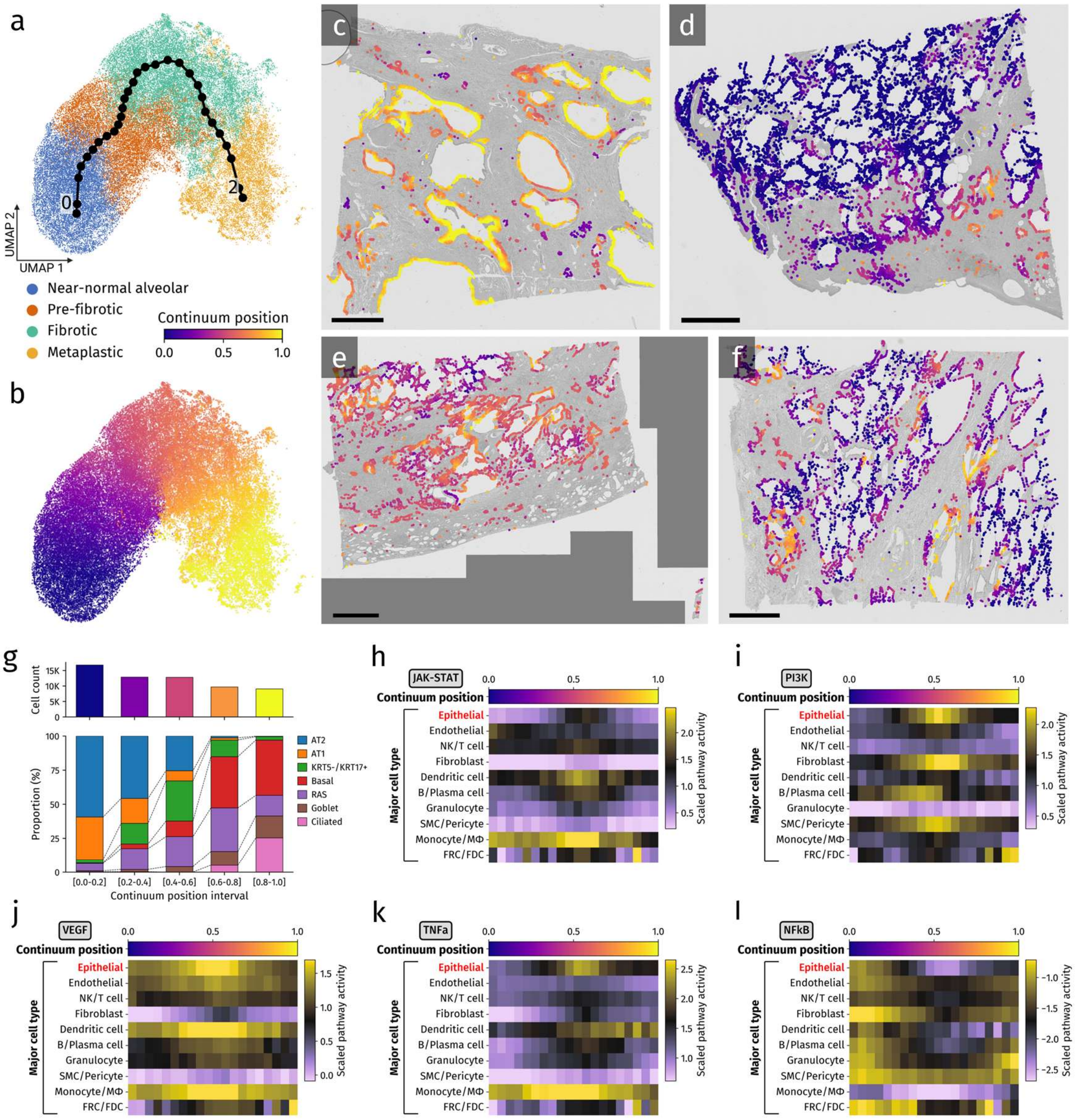


**Extended Data Fig. 7 | Pseudo-temporal continuum inference and dynamics of epithelial and microenvironmental pathway activity along the continuum.**

**a,** UMAP embedding of epithelial-centered neighborhood profiles with the inferred trajectory from scFates. Bronchiolar, alveolar duct, and epithelial-rich neighborhood classes were excluded from this analysis to focus on fibrotic remodeling rather than normal airway-associated structures or epithelial-dominant profiles with limited non-epithelial neighborhood information. **b,** UMAP embedding of epithelial-centered neighborhood profiles color-coded by continuum position. **c–f,** Spatial maps of epithelial cells color-coded by continuum position overlaid on grayscale tissue images in HC_s1 (c), AF_s2 (d), FR_s3 (e), and FR_s4 (f). Scale bars, 1,000 μm. **g,** Bar plots showing the number of epithelial cells in each continuum-position interval (top) and stacked bar plots showing the epithelial subtype composition within each interval (bottom). **h–l,** Heatmaps showing JAK–STAT (h), PI3K (i), VEGF (j), TNFα (k), and NF-κB (l) pathway activity in epithelial and nearby non-epithelial major cell types along the continuum. Activity scores were averaged within continuum-position bins and scaled for visualization.

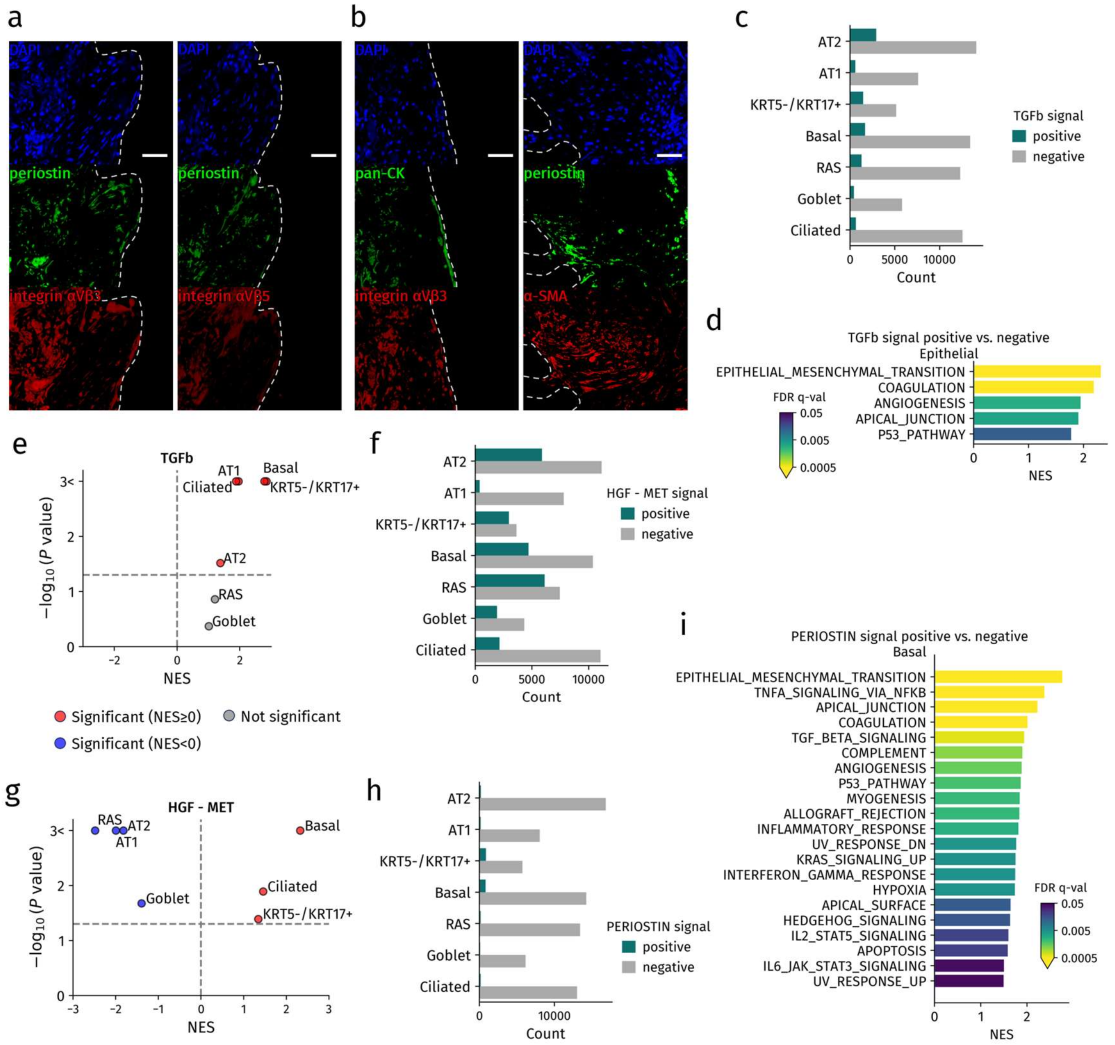


**Extended Data Fig. 8 | Periostin-family signaling in fibroblastic foci and validation of the cell-level positive-versus-negative comparison strategy. a,** Representative immunofluorescence images of fibroblastic foci in serial sections from FR_s3, showing DAPI (blue), periostin (green), and integrin αVβ3 or integrin αVβ5 (red). Scale bars, 50 µm. **b,** Representative immunofluorescence images of fibroblastic foci in serial sections from FR_s3, showing pan-CK, periostin, integrin αVβ3, and α-SMA. Co-staining showed periostin in α-SMA-positive myofibroblast-rich regions and integrin αVβ3 in pan-CK-positive epithelium overlying fibroblastic foci. Scale bars, 50 µm. **c,** Bar plots showing the number of signal-positive and signal-negative epithelial cells for incoming

TGFβ-family interactions from neighboring cells within each epithelial subtype. **d,** Gene set enrichment analysis of basal cells comparing cells positive versus negative for inferred incoming TGFβ-family signals from neighboring cells. Bars show normalized enrichment scores (NES) for significantly enriched Hallmark gene sets (FDR q value < 0.05), with color indicating the FDR q value. **e,** Plot showing the NES and P value of the epithelial–mesenchymal transition Hallmark gene set for comparisons between cells positive versus negative for inferred incoming TGFβ-family signals from neighboring cells across epithelial subtypes. Red indicates significantly positive enrichment, and gray indicates not significant. **f,** Bar plots showing the numbers of signal-positive and signal-negative epithelial cells for HGF–MET interaction from neighboring cells within each epithelial subtype. **g,** Plot showing the NES and P value of the epithelial–mesenchymal transition Hallmark gene set for comparisons between cells positive versus negative for inferred incoming HGF–MET signal from neighboring cells across epithelial subtypes. Red indicates significantly positive enrichment and blue indicates significantly negative enrichment. **h,** Bar plots showing the number of signal-positive and signal-negative epithelial cells for incoming PERIOSTIN-family interactions from neighboring cells within each epithelial subtype. **i,** Gene set enrichment analysis of basal cells comparing cells positive versus negative for inferred incoming PERIOSTIN-family signals from neighboring cells. Bars show NES for significantly enriched Hallmark gene sets (FDR q value <0.05), with color indicating the FDR q value.

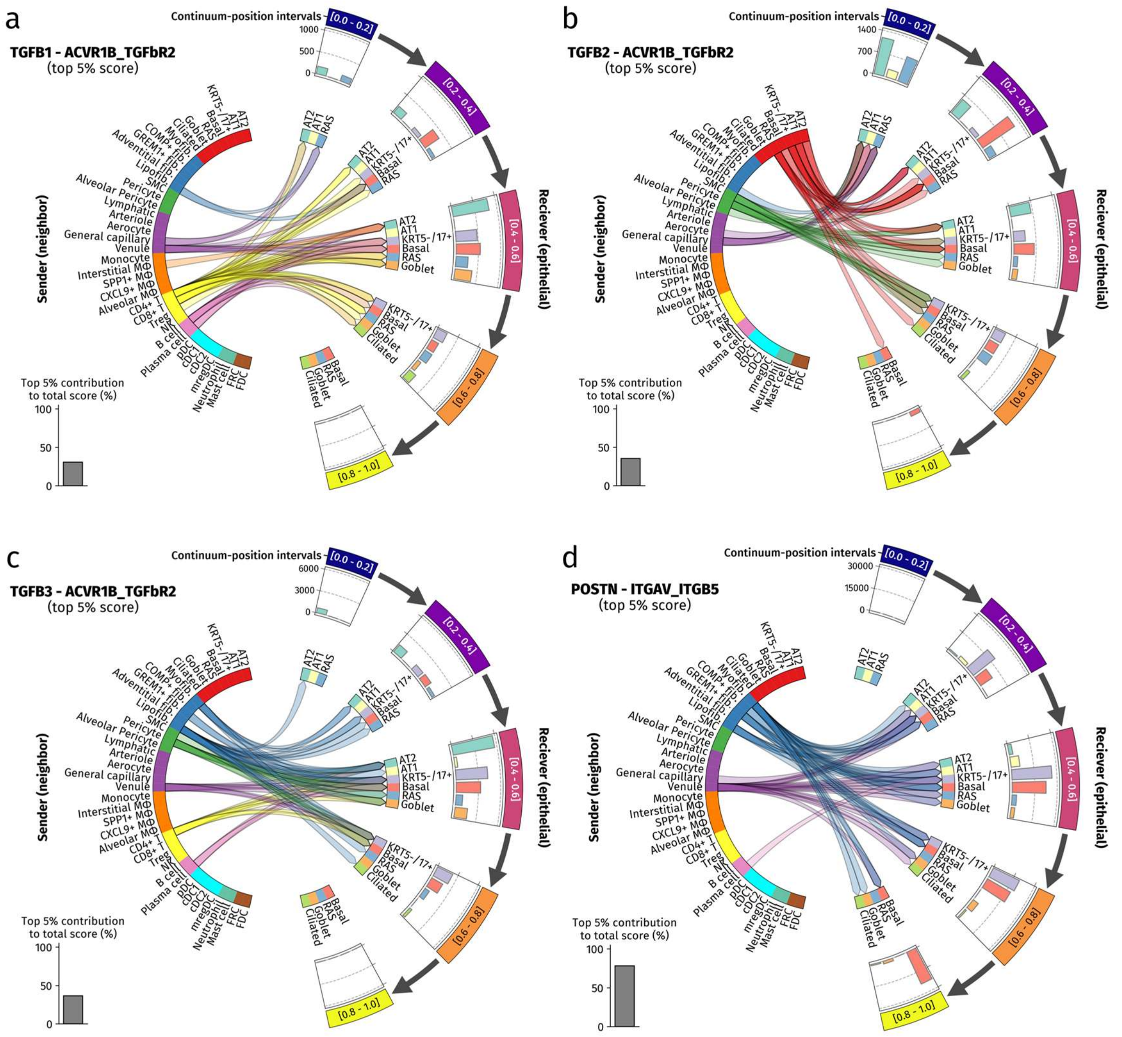


**Extended Data Fig. 9 | Dynamic changes in cell–cell interactions along the pseudo-temporal continuum from neighboring cells to epithelial subtypes. a–d,** Circular plots showing the distribution of representative interactions across cell subtypes and continuum-position intervals for interactions from neighboring cells to epithelial cells: TGFB1–ACVR1B_TGFbR2 (a), TGFB2–ACVR1B_TGFbR2 (b), TGFB3–ACVR1B_TGFbR2 (c), and POSTN–ITGAV_ITGB5 (d). Epithelial cells were grouped into five continuum-position intervals. Epithelial subtype–interval combinations in which the subtype accounted for less than 3% of epithelial cells in that interval were excluded from visualization. Links indicate interactions between sender and receiver cell subtypes, and only interactions belonging to the top 5% of scores are shown. Bar plots above the epithelial sectors show the summed interaction score within each continuum-position interval. The inset bar plot indicates the percentage contribution of the top 5% of interaction scores to the total score.

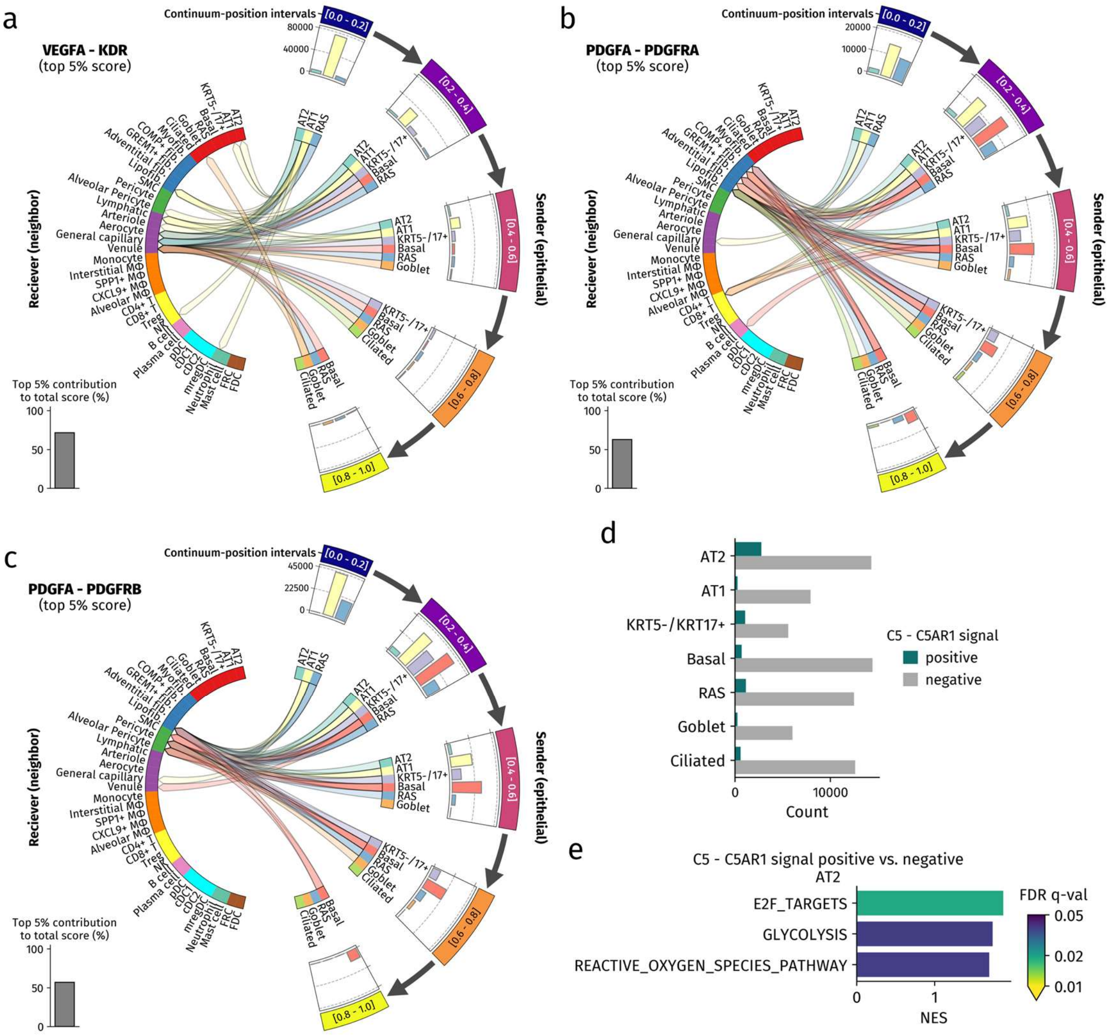


**Extended Data Fig. 10 | Dynamic changes in cell–cell interactions along the pseudo-temporal continuum from epithelial subtypes to neighboring cells. a–c,** Circular plots showing the distribution of representative interactions across cell subtypes and continuum-position intervals for interactions from epithelial cells to neighboring cells: VEGFA–KDR (a), PDGFA–PDGFRA (b), and PDGFA–PDGFRB (c). Epithelial cells were grouped into five continuum-position intervals. Epithelial subtype–interval combinations in which the subtype accounted for less than 3% of epithelial cells in that interval were excluded from visualization. Links indicate interactions between sender and receiver cell subtypes, and only interactions belonging to the top 5% of scores are shown. Bar plots above the epithelial sectors show the summed interaction score within each continuum-position interval. The inset bar plot indicates the percentage contribution of the top 5% of interaction scores to the total score. **d,** Bar plots showing the number of signal-positive and signal-negative cells in each epithelial subtype for the outgoing C5–C5AR1 interaction from epithelial cells to neighboring cells. **e,** Gene set enrichment analysis of AT2 cells comparing cells positive versus negative for the inferred outgoing C5–C5AR1 interaction signal. Bars show normalized enrichment scores (NES) for significantly enriched Hallmark gene sets (FDR q value <0.05), with color indicating the FDR q value.

#### Supplementary Figure Legends

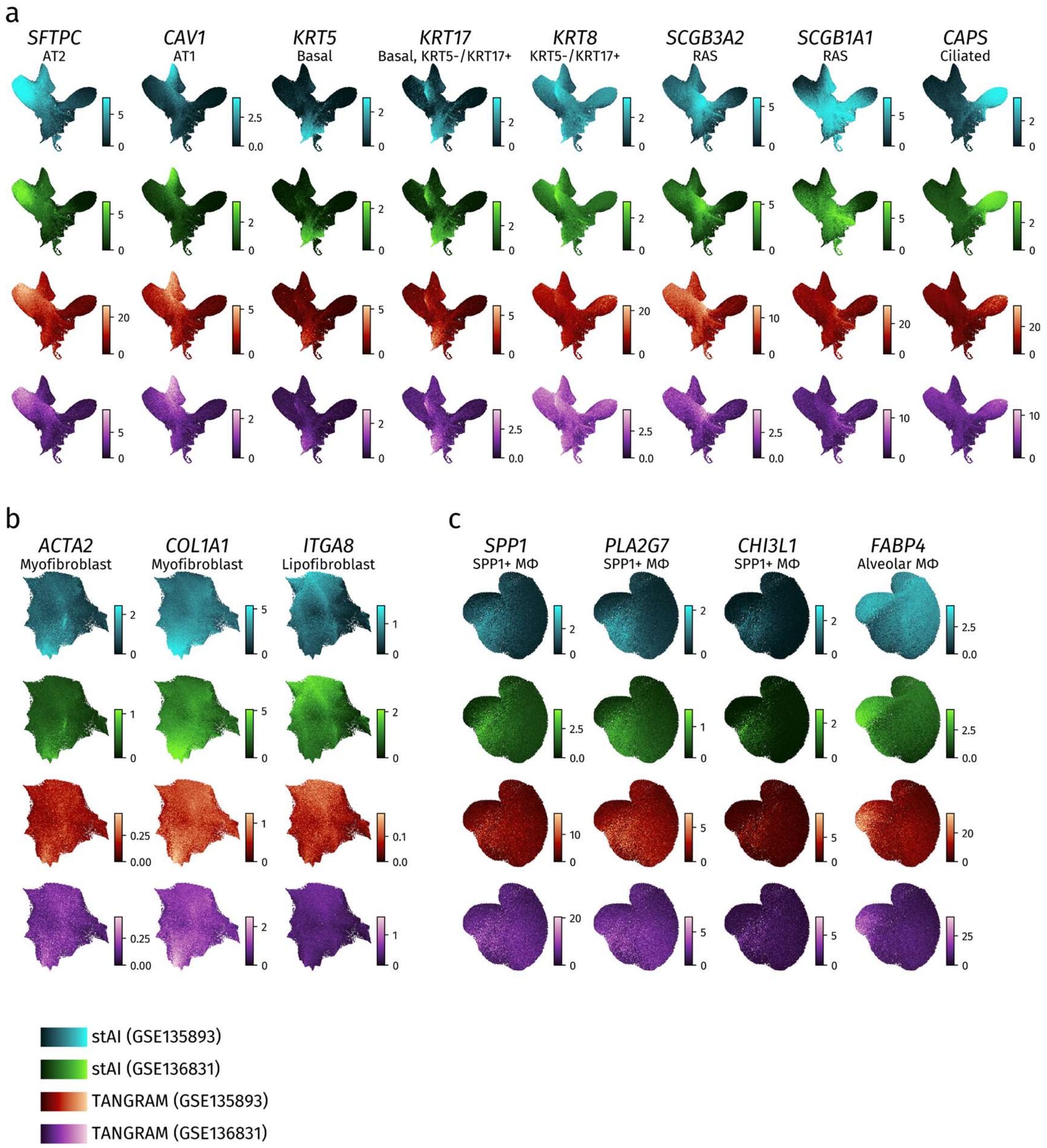


**Supplementary Fig. 1| Imputed marker gene expression supporting subtype annotation. a–c,** Feature plots showing imputed expression of marker genes projected onto UMAP embeddings of epithelial cells (a), fibroblasts (b), and macrophages (c). Imputation was performed using stAI or Tangram with GSE135894 and GSE 136831 as reference datasets. Rows indicate the imputation method and reference dataset, and columns indicate marker genes corresponding to the indicated subtypes. Color intensity indicates imputed expression level.

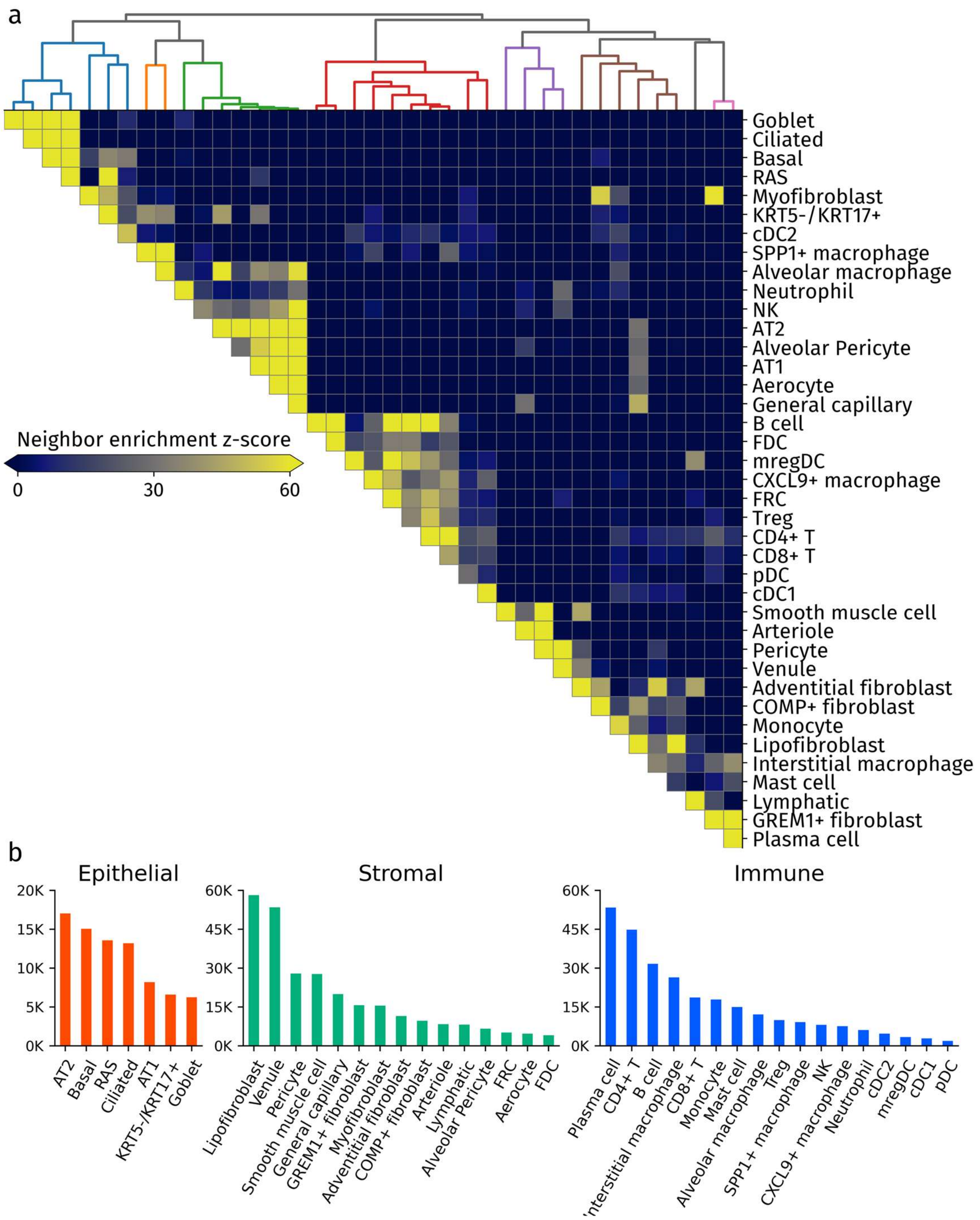


**Supplementary Fig. 2 | Global neighborhood enrichment and cell subtype counts across samples. a,** Heatmap showing pairwise neighborhood enrichment z-scores among annotated cell subtypes calculated using Squidpy. Cell subtypes were ordered according to similarity in their neighborhood enrichment patterns, as shown by the dendrogram. Color intensity indicates the neighborhood enrichment z-score. **b,** Bar plots showing total cell counts for each annotated subtype across all six samples, grouped into epithelial, stromal, and immune compartments; K denotes thousands.

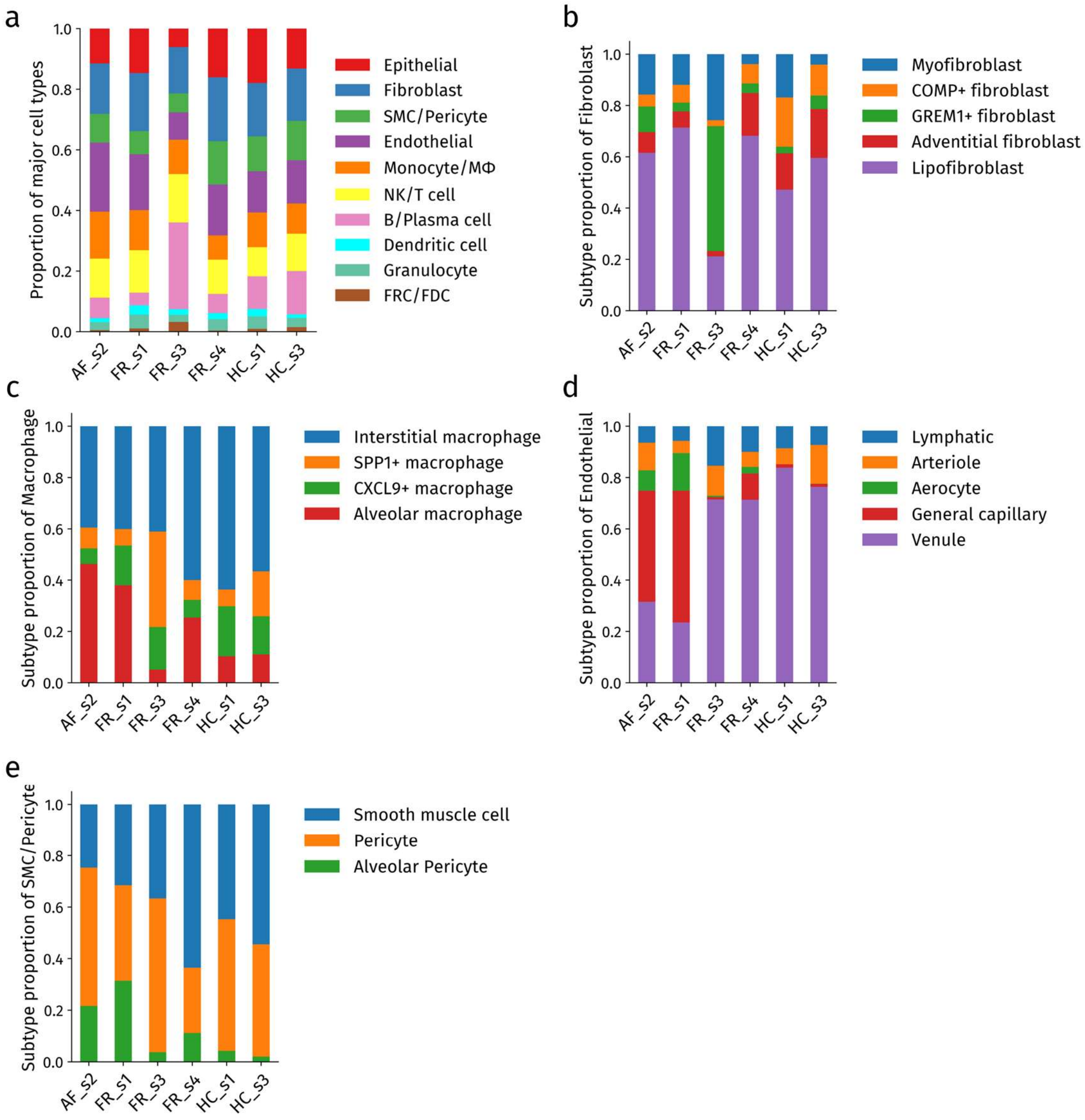


**Supplementary Fig. 3 | Major cell-type and selected lineage-specific subtype compositions across samples. a,** Stacked bar plots showing the major cell-type composition in each sample. **b–e,** Stacked bar plots showing subtype composition within fibroblasts (b), macrophages (c), endothelial cells (d) and smooth muscle cells/pericytes (e) in each sample. Values represent proportions among all cells (a) or within the indicated lineage (b–e). FRC, fibroblastic reticular cell; FDC, follicular dendritic cell; Mφ, macrophage.

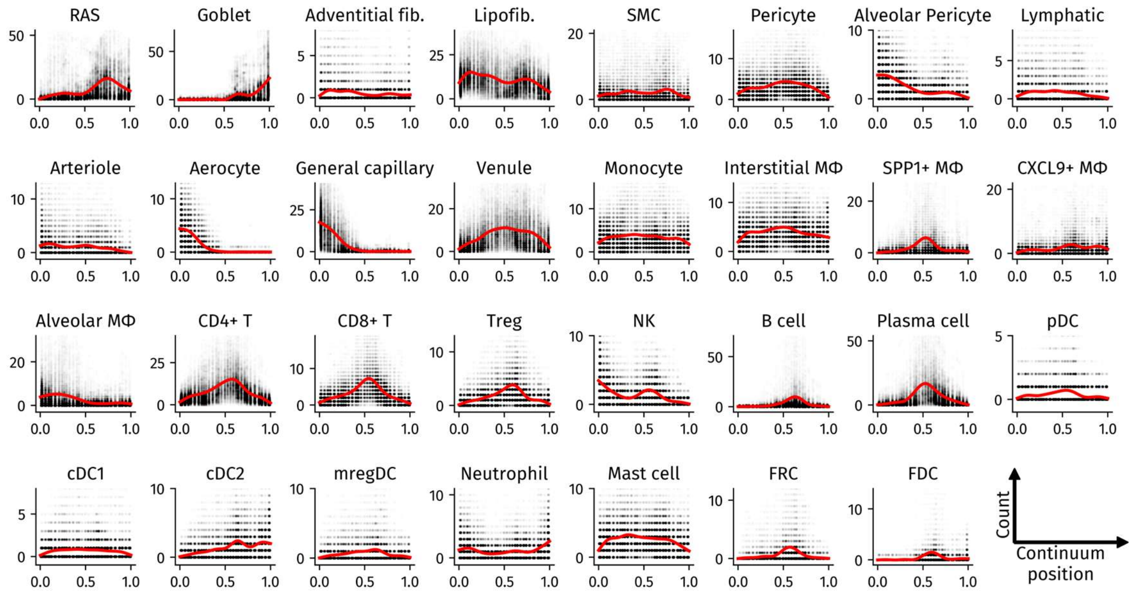


**Supplementary Fig. 4 | Changes in the numbers of subtypes within epithelial-centered neighborhood profiles along the continuum.** Changes in the numbers of remaining annotated subtypes, complementing the selected epithelial and fibroblast subtypes shown in Fig. 3f, g. Gray dots indicate observed neighborhood counts, and red curves indicate fitted trends inferred by scFates.

#### Supplementary Tables

##### Supplementary Table 1: Patient characteristics and sample information

| Patient ID | s1 | s2 | s3 | s4 |
|---|---|---|---|---|
| Sample ID | FR_s1<br>HC_s1 | AF_s2 | FR_s3<br>HC_s3 | FR_s4 |
| Age | 86 | 72 | 68 | 82 |
| Sex | Male | Male | Male | Male |
| Smoking history (pack-years) | 40 | 150 | 45 | 30 |
| Resection site | Left upper lobe (wedge resection) | Left upper lobe (wedge resection) | Left upper lobe (wedge resection) | Right lower lobe (segmentectomy) |
| Histopathological pattern | UIP | UIP | probable UIP | UIP |

AF, alveolar–fibrotic interface region; FR, fibroblastic focus-rich region; HC, honeycombing-dominant region; UIP, usual interstitial pneumonia.

**Supplementary Table 3: Marker genes supporting cell-type and subtype annotation**

| Major cell type | Cell subtype | Marker genes | Imputed markers for validation |
|---|---|---|---|
| Epithelial | | *EPCAM, CDH1* | |
| | AT2 | *LAMP3, ABCA3* | *SFTPC* |
| | AT1 | *AGER, RTKN2* | *CAV1* |
| | *KRT5*$^-$/*KRT17*$^+$ | *ITGB6, CDKN2B* | *KRT8, KRT17* |
| | Basal | *ANXA8, TP63* | *KRT5, KRT17* |
| | RAS | | *SCGB3A2, SCGB1A1* |
| | Goblet | *MUC5B, FCGBP* | |
| | Ciliated | *FOXJ1, TPPP3* | *CAPS* |
| Fibroblast | | *COL5A1, PDGFRA* | |
| | Myofibroblast | *POSTN, CTHRC1* | *ACTA2, COL1A1* |
| | *COMP*$^+$ fibroblast | *COMP* | |
| | *GREM1*$^+$ fibroblast | *GREM1* | |
| | Lipofibroblast | *TCF21, AOC3* | *ITGA8* |
| | Adventitial fibroblast | *PI16, CD34* | |
| Monocyte/MΦ | | *CD14, SLCO2B1* | |
| | Monocyte | *FCN1* | |
| | Interstitial macrophage | *F13A1* | |
| | *SPP1*$^+$ macrophage | *MMP9* | *SPP1, PLA2G7, CHI3L1* |
| | *CXCL9*$^+$ macrophage | *CXCL9* | |
| | Alveolar macrophage | *SLCO2B1, ALDH2* | *FABP4* |
| SMC/Pericyte | | | |
| | Smooth muscle cell | *MYLK, CSRP1* | |
| | Pericyte | *PDGFRB, STEAP4* | |
| | Alveolar Pericyte | *PDGFRB, KCNK3* | |
| Endothelial | | *CLDN5, SHANK3* | |
| | Lymphatic | *PROX1* | |
| | Arteriole | *GJA5, VEGFC* | |
| | Aerocyte | *HPGD, TBX2* | |
| | General capillary | *SLC6A4, TMEM100* | |
| | Venule | *PLVAP* | |
| NK/T cell | | | |
| | CD4$^+$ T | *CD3E*, *CD8A*$^-$, *FOXP3*$^-$ | |
| | CD8$^+$ T | *CD3E, CD8A, GZMH* | |
| | Treg | *CD3E, FOXP3* | |
| | NK | *PRF1, GZMH* | |
| B/Plasma | | | |

| | | |
|---|---|---|
| | B cell | *MS4A1, CD19, CR2* |
| | Plasma cell | *XBP1, DERL3* |
| Dendritic cell | | |
| | pDC | *LILRA4, IRF7* |
| | cDC1 | *WDFY4, CLEC9A* |
| | cDC2 | *CD1C, FCER1A* |
| | mregDC | *CCL22, FSCN1* |
| Granulocyte | | |
| | Neutrophil | *CXCR1, DEFA1* |
| | Mast cell | *KIT, GATA2* |
| Others | | |
| | FDC | *CXCL13, CR2* |
| | FRC | *CCL19* |

Marker genes were used to support annotation. Imputed markers were assessed after expression imputation for annotation validation. RAS, respiratory airway secretory cells. pDC, plasmacytoid dendritic cells. cDC, conventional dendritic cells. mregDC, mature dendritic cells enriched in immunoregulatory molecules. FDC, follicular dendritic cells. FRC; fibroblastic reticular cells.

**Supplementary Table 10: Cell-type composition by samples (%)**

| Cell type | AF_s2 | FR_s1 | FR_s3 | FR_s4 | HC_s1 | HC_s3 |
|---|---|---|---|---|---|---|
| AT2 | 6.84 | 5.24 | 0.93 | 3.61 | 0.21 | 1.30 |
| AT1 | 2.63 | 3.02 | 0.30 | 2.08 | 0.14 | 0.63 |
| *KRT5*$^-$/*KRT17*$^+$ | 0.60 | 1.64 | 1.46 | 1.01 | 0.30 | 1.14 |
| Basal | 0.49 | 0.56 | 1.12 | 2.49 | 6.65 | 2.85 |
| RAS | 0.69 | 2.58 | 1.39 | 2.87 | 2.56 | 3.32 |
| Goblet | 0.15 | 0.29 | 0.56 | 1.11 | 2.87 | 0.77 |
| Ciliated | 0.11 | 1.32 | 0.37 | 2.86 | 5.23 | 3.13 |
| Myofibroblast | 2.61 | 2.30 | 3.92 | 0.82 | 2.96 | 0.70 |
| *COMP*$^+$ fibroblast | 0.78 | 1.33 | 0.36 | 1.59 | 3.38 | 2.09 |
| *GREM1*$^+$ fibroblast | 1.66 | 0.66 | 7.42 | 0.79 | 0.44 | 0.91 |
| Adventitial fibroblast | 1.34 | 1.22 | 0.31 | 3.52 | 2.49 | 3.30 |
| Lipofibroblast | 10.22 | 13.70 | 3.23 | 14.40 | 8.29 | 10.33 |
| Smooth muscle cell | 2.35 | 2.41 | 2.29 | 9.08 | 5.17 | 7.07 |
| Pericyte | 5.11 | 2.82 | 3.73 | 3.62 | 5.89 | 5.65 |
| Alveolar Pericyte | 2.06 | 2.39 | 0.23 | 1.60 | 0.49 | 0.26 |
| Lymphatic | 1.46 | 1.05 | 1.40 | 1.68 | 1.16 | 1.04 |
| Arteriole | 2.45 | 0.89 | 1.06 | 0.99 | 0.85 | 2.16 |
| Aerocyte | 1.82 | 2.70 | 0.05 | 0.43 | 0.00 | 0.01 |
| General capillary | 9.85 | 9.49 | 0.08 | 1.72 | 0.18 | 0.16 |
| Venule | 7.18 | 4.33 | 6.48 | 11.98 | 11.39 | 10.91 |
| Monocyte | 3.26 | 4.20 | 1.86 | 1.75 | 4.12 | 1.85 |
| Interstitial macrophage | 4.84 | 3.60 | 3.89 | 3.73 | 4.67 | 4.56 |
| SPP1+ macrophage | 1.00 | 0.57 | 3.52 | 0.48 | 0.48 | 1.41 |
| *CXCL9*$^+$ macrophage | 0.74 | 1.39 | 1.57 | 0.43 | 1.43 | 1.20 |
| Alveolar macrophage | 5.68 | 3.41 | 0.49 | 1.58 | 0.76 | 0.89 |
| CD4$^+$ T | 6.25 | 8.10 | 8.43 | 6.33 | 5.96 | 6.78 |
| CD8$^+$ T | 2.83 | 2.80 | 3.95 | 2.88 | 1.97 | 2.89 |
| Treg | 1.61 | 1.52 | 2.04 | 1.26 | 1.11 | 1.69 |
| NK | 2.18 | 1.57 | 1.51 | 0.86 | 0.56 | 1.02 |
| B cell | 3.80 | 1.91 | 7.89 | 2.44 | 5.71 | 6.17 |
| Plasma cell | 2.91 | 2.27 | 20.69 | 3.83 | 5.09 | 8.06 |
| pDC | 0.22 | 0.53 | 0.38 | 0.23 | 0.22 | 0.18 |
| cDC1 | 0.39 | 1.00 | 0.28 | 0.39 | 0.48 | 0.22 |
| cDC2 | 0.29 | 0.69 | 0.62 | 0.95 | 1.35 | 0.49 |
| mregDC | 0.47 | 0.92 | 0.61 | 0.52 | 0.37 | 0.31 |
| Neutrophil | 0.79 | 1.63 | 0.73 | 0.71 | 1.09 | 0.92 |
| Mast cell | 1.87 | 2.94 | 1.63 | 3.02 | 3.02 | 2.15 |
| FRC | 0.22 | 0.83 | 1.57 | 0.21 | 0.55 | 0.98 |
| FDC | 0.27 | 0.20 | 1.64 | 0.16 | 0.42 | 0.51 |

Values indicate the percentage of each cell type among all cells passing quality control in each sample.
