## Supplementary Note for "Trajectory inference of epithelial-centered neighborhood profiles reconstructs a pseudo-temporal continuum in idiopathic pulmonary fibrosis"

**Spatial distance-weighted ligand–receptor interaction scoring**

We quantified distance-weighted ligand–receptor–mediated interaction scores between epithelial cells and their spatial neighbors using a curated interaction database and spatially resolved transcriptomes. The raw count matrix was first normalized on a per-cell basis to a constant library size, yielding normalized expression values $x_{cg}$ for cell $c$ and gene $g$. Ligand–receptor interactions were imported from the CellChat database[9] and restricted to those annotated as "Secreted Signaling." For each interaction $p$, we defined the ligand gene set $L_p$ and receptor gene set $R_p$, and their union $G_p = L_p \cup R_p$. We then intersected the union of all $G_p$ with the genes represented in the Xenium gene panel and retained only those interactions for which all constituent genes were present in the expression matrix ($G_p \subseteq \mathcal{G}$), where $\mathcal{G}$ denotes the set of genes in the dataset. The resulting interaction table was provided as a supplementary resource (Supplementary Table 10). To summarize ligand and receptor activities, we used the geometric mean of normalized expression values without adding a per-gene rescaling or pseudocount addition. For any gene set $G \subseteq \mathcal{G}$, its activity in cell $c$ was defined as

$$\mathrm{GM}_c(G) = \begin{cases} \exp\left(\frac{1}{\mid G \mid}\sum_{g \in G} \log x_{cg}\right), & \text{if } x_{cg} > 0 \text{ for all } g \in G \\ 0, & \text{otherwise.} \end{cases}$$

Thus, the activity of a ligand or receptor complex was set to zero if any constituent gene showed zero normalized expression in that cell. For each interaction $p$, ligand and receptor activities in cell $c$were then defined as

$$L_{cp} = \mathrm{GM}_c(L_p), \;\; R_{cp} = \mathrm{GM}_c(R_p).$$

For each cell $c$, let $\mathbf{s}_c \in \mathbb{R}^2$ denote its two-dimensional spatial coordinates. To enable unified processing across samples, cells from all samples were embedded in a shared coordinate space after applying sample-specific spatial offsets based on the spatial extent of each sample, such that the distance between any two cells from different samples exceeded the neighborhood radius. Epithelial cells were used as focal cells. For each epithelial cell $i \in \mathcal{C}_{epi}$, we defined its local spatial neighborhood as

$$\mathcal{N}(i) = \{ j \in \mathcal{C} \mid \ \| \mathbf{s}_j - \mathbf{s}_i \| \leq r, j \neq i \},$$

where $\mathcal{C}$ denotes the set of all cells across samples and $r = 300\ \mu$m. Since all inter-sample cell-to-cell distances exceeded $r$ after offsetting, only within-sample neighbors could contribute to the interaction scores. Radius-based neighbor searches were implemented using SciPy's cKDTree[10]. To model the decay of interaction strength with distance, we assigned each epithelial–neighbor pair $(i, j)$ a Gaussian distance weight

$$w_{ij} = \exp\left(-\frac{d_{ij}^2}{2\sigma^2}\right),\ d_{ij} = \| \mathbf{s}_j - \mathbf{s}_i \|,$$

with $\sigma = 150\ \mu$m. Thus, cells closer to the epithelial cell contributed more strongly than more distant neighbors. These weights were precomputed for all epithelial-neighbor pairs in the distance-weighted neighborhood graph. For each interaction $p$ and each epithelial–neighbor pair $(i, j)$, we computed directional ligand–receptor interaction scores in both neighbor-to-epithelial and epithelial-to-neighbor directions as,

$$S_{j \to i}^{(p)} = L_{jp}\, R_{ip}\, w_{ij},\ S_{i \to j}^{(p)} = L_{ip}\, R_{jp}\, w_{ij}.$$

Here, $S_{j \to i}^{(p)}$ represents a putative incoming signal from neighboring cell $j$ to epithelial cell $i$ via interaction $p$, whereas $S_{i \to j}^{(p)}$ represents the corresponding outgoing signal from epithelial cell $i$to neighboring cell $j$.

**Aggregation of cell–cell interaction by subtype**

To assess the significance of distance-weighted ligand–receptor interaction scores at the subtype level, we aggregated epithelial centered cell–cell edges by subtype and performed permutation-based testing. For each interaction $p$ and each direction, we constructed a subtype-by-subtype interaction matrix with epithelial subtypes on one axis and dataset-wide cell subtypes on the other. For incoming signals (neighboring-cell-to-epithelial), rows represented neighboring-cell subtypes and columns represented epithelial subtypes. For outgoing signals (epithelial-to-neighboring-cell), rows represented epithelial subtypes and columns represented neighboring-cell subtypes. Each epithelial–neighbor edge carried directional interaction scores $S_{j\to i}^{(p)}$ and $S_{i\to j}^{(p)}$, where $i$ denotes an epithelial cell and $j$ denotes one of its neighboring cells. To account for differences in the abundance of epithelial cells across epithelial subtypes, edge scores were normalized by the number of epithelial cells in the relevant epithelial subtype. Specifically, for incoming signals from neighboring subtype $s$ to epithelial subtype $t$, the observed aggregated score was defined as

$$M_{n\to e}^{\mathrm{obs},(p)}(s,t) = \frac{1}{N_t^{\mathrm{epi}}} \sum_{(j,i)\in E_{st}^{n\to e}} S_{j\to i}^{(p)},$$

where $E_{st}^{n\to e}$ denotes the set of edges from neighboring cells of subtype $s$to epithelial cells of subtype $t$, and $N_t^{\mathrm{epi}}$ denotes the number of epithelial cells in subtype $t$. Likewise, for outgoing signals from epithelial subtype $s$ to neighboring subtype $t$, the observed aggregated score was defined as

$$M_{e\to n}^{\mathrm{obs},(p)}(s,t) = \frac{1}{N_s^{\mathrm{epi}}} \sum_{(i,j)\in E_{st}^{e\to n}} S_{i\to j}^{(p)},$$

where $E_{st}^{e\to n}$ denotes the set of edges from epithelial cells of subtype $s$ to neighboring cells of subtype $t$, and $N_s^{\mathrm{epi}}$ denotes the number of epithelial cells in subtype $s$. To evaluate whether the observed subtype-level interaction scores exceeded random expectation, neighboring-cell subtype labels were permuted across cells while epithelial subtype labels were kept fixed. This permutation scheme was applied to both directions, such that the neighboring-cell subtype labels associated with each edge were shuffled for both neighboring-cell-to-epithelial and epithelial-to-neighboring-cell interaction matrices. In each of $B = 10{,}000$ iterations, neighboring-cell subtype labels were randomly reassigned and the corresponding permuted matrices, $M_{n\to e}^{\mathrm{perm},b,(p)}(s,t)$ and $M_{e\to n}^{\mathrm{perm},b,(p)}(s,t)$, were recalculated. For each subtype pair and interaction, we counted the number of permutation replicates in which the permuted score was greater than or equal to the observed score:

$$C_{n\to e}^{(p)}(s,t) = \sum_{b=1}^{B} \mathbb{1}\left[M_{n\to e}^{\mathrm{perm},b,(p)}(s,t) \geq M_{n\to e}^{\mathrm{obs},(p)}(s,t)\right],$$

$$C_{e\to n}^{(p)}(s,t) = \sum_{b=1}^{B} \mathbb{1}\left[M_{e\to n}^{\mathrm{perm},b,(p)}(s,t) \geq M_{e\to n}^{\mathrm{obs},(p)}(s,t)\right].$$

Permutation $P$-values were estimated using a +1 correction to avoid zero counts:

$$P^{(p)}(s,t) = \frac{C^{(p)}(s,t) + 1}{B + 1}.$$

Multiple testing correction across subtype pairs was performed using the Benjamini-Hochberg procedure with a false discovery rate threshold of 0.05. This procedure produced a summary table reporting, for each interaction, direction, and subtype pair, the observed normalized score, the number of permutation replicates with scores at least as large as the observed value, the total number of permutations, the permutation $P$-value, and the Benjamini-Hochberg-adjusted $P$-value (FDR) (Supplementary Table 11, 12). For visualization, interactions declared significant at FDR <0.05 were ranked using log-transformed and min-max normalized observed scores. For each direction, the top 50 subtype pairs were used to construct Sankey diagrams, with source and target subtypes shown as distinct node sets. Link widths were proportional to the raw observed scores, and link colors were matched to the sender subtype with semi-transparency to indicate flow direction. To summarize interaction-specific patterns across multiple source-to-target pairs, the top 50 interactions were further arranged into a matrix with rows representing source-to-target subtype pairs and columns representing interaction labels, and nonzero entries were displayed as scatter points colored by normalized score. A horizontal color bar indicated the normalized signal scale from 0 to 1.
